# An endogenous GLP-1 circuit engages VTA GABA neurons to regulate mesolimbic dopamine neurons and attenuate cocaine seeking

**DOI:** 10.1101/2024.06.20.599574

**Authors:** Riley Merkel, Nicole Hernandez, Vanessa Weir, Yafang Zhang, Matthew T. Rich, Richard C. Crist, Benjamin C. Reiner, Heath D. Schmidt

**Author notes:** Corresponding Author: Heath D. Schmidt, Ph.D., Professor & Killebrew-Censits Chair of Undergraduate Education Department of Biobehavioral Health Sciences, School of Nursing Department of Psychiatry, Perelman School of Medicine University of Pennsylvania, 125 South 31^st^ Street, TRL Room 2215 Philadelphia, PA, 19104.

## Abstract

Recent studies show that systemic administration of a glucagon-like peptide-1 receptor (GLP-1R) agonist is sufficient to attenuate the reinstatement of cocaine-seeking behavior, an animal model of relapse. However, the neural mechanisms mediating these effects and the role of endogenous central GLP-1 signaling in cocaine seeking remain unknown. Here, we show that voluntary cocaine taking decreased plasma GLP-1 levels in rats and that chemogenetic activation of GLP-1-producing neurons in the nucleus tractus solitarius (NTS) that project to the ventral tegmental area (VTA) decreased cocaine reinstatement. Single nuclei transcriptomics and FISH studies revealed GLP-1Rs are expressed primarily on GABA neurons in the VTA. Using *in vivo* fiber photometry, we found that the efficacy of a systemic GLP-1R agonist to attenuate cocaine seeking was associated with increased activity of VTA GABA neurons and decreased activity of VTA dopamine neurons. Together, these findings suggest that targeting central GLP-1 circuits may be an effective strategy toward reducing cocaine relapse and highlight a novel functional role of GABAergic GLP-1R-expressing midbrain neurons in drug seeking.

## Introduction

Cocaine use disorder (CUD) remains a significant public health concern in the United States. Unfortunately, the prevalence of cocaine use, as well as the incidence of fatal overdoses involving cocaine, have increased year-over-year^1,2^. Despite decades of focused pre-clinical and clinical studies that have advanced our understanding of the anatomical, neurochemical, molecular, and epigenetic bases of psychostimulant addiction, a safe and efficacious pharmacotherapy for CUD remains to be discovered^3^. Thus, an improved understanding of the neurobiological mechanisms that regulate cocaine taking and seeking is needed to inform conceptually new approaches to treating CUD.

An emerging literature indicates that activation of central glucagon-like peptide-1 receptors (GLP-1Rs) reduces the rewarding and reinforcing effects of addictive drugs^4,5,6,7^. Previously, we identified a systemic dose of the GLP-1R agonist exendin-4 that selectively attenuated the reinstatement of cocaine-seeking behavior in rats^8^, effects mediated, in part, by activation of GLP-1Rs in the ventral tegmental area (VTA)^8^. Consistent with these effects, direct activation of GLP-1Rs in the VTA is sufficient to attenuate cocaine seeking^8^. In addition to expressing GLP-1Rs^9,10^, the VTA also receives direct monosynaptic projections from GLP-1-producing preproglucagon (PPG) neurons in the nucleus tractus solitarius (NTS)^11,12^. Together with our behavioral pharmacology studies, this anatomy suggests that activating endogenous GLP-1-producing NTS→VTA projections may suppress cocaine-seeking behaviors during abstinence. However, no studies, to date, have targeted endogenous GLP-1-producing NTS circuits to reduce drug-seeking behavior.

The mechanisms by which GLP-1R activation in the midbrain attenuates cocaine seeking remain unknown. Therefore, phenotyping GLP-1R-expressing cells in the VTA and determining how GLP-1R activation alters activity of these neurons during cocaine seeking are essential ‘next steps’ toward developing the next-generation of GLP-1R-based therapies to treat CUD. In the present study, we: 1) assessed the functional role of endogenous VTA GLP-1 signaling in cocaine seeking using projection-specific chemogenetic approaches, 2) determined cell type-specific expression patterns of GLP-1Rs in the VTA using fluorescent *in situ* hybridization (FISH) and single nuclei RNA-sequencing (snRNA-seq) methods, and 3) used *in vivo* fiber photometry in freely moving transgenic rats to further our understanding of the cell type-specific mechanisms in the midbrain underlying the suppressive effects of a GLP-1R agonist on cocaine seeking. Our findings indicate that voluntary cocaine taking decreased circulating GLP-1 levels and that chemogenetic activation of NTS→VTA projections was sufficient to attenuate cocaine seeking via increased GLP-1 signaling in the VTA. Using FISH and snRNA-seq, we phenotyped GLP-1R-expressing cells in the midbrain and discovered that GLP-1Rs are expressed primarily on GABA neurons in the VTA. Lastly, using *in vivo* calcium imaging we showed that the efficacy of systemic exendin-4 is associated with dynamic changes in activity of VTA GABA and dopamine neurons that are associated with decreased cocaine seeking. Together, these findings reveal a functionally relevant cell type-specific mechanism underlying the suppressive effects of GLP-1R agonists on cocaine seeking and highlight an endogenous GLP-1-expressing circuit that could be targeted to treat CUD and reduce cocaine relapse.

## Methods

### Animals and housing

Male Sprague-Dawley rats (*Rattus norvegicus*) weighing 225-250g were obtained from Taconic Laboratories (Rensselaer, NY). Male and female transgenic Long-Evans rats (*Rattus norvegicus*) expressing Cre recombinase under the rat *Gad1* promoter (LE-Tg(Gad1-iCre)3Ottc) were purchased from the Rat Resource and Research Center (RRRC P40OD011062; Columbia, MO)^13,14,15^. Male and female transgenic Sprague-Dawley rats (*Rattus norvegicus)* expressing Cre recombinase under the rat *Th* promoter (HsdSage:SD-*TH^em1(IRES-Cre)Sage^*) were purchased from Envigo (Indianapolis, IN)^16,17^. Rats were housed individually on a 12/12 h light/dark cycle and maintained on *ad libitum* food and water. All experimental procedures were performed during the light cycle. The experimental protocols were consistent with the guidelines issued by the U.S. National Institutes of Health and were approved by the University of Pennsylvania’s Institutional Animal Care and Use Committee.

### Drugs

Cocaine HCl was obtained from the National Institute on Drug Abuse (Rockville, MD) and dissolved in bacteriostatic 0.9% saline. Exendin-(9-39) and exendin-4 were purchased from Bachem (Torrance, CA) and dissolved in artificial cerebrospinal fluid (aCSF; Harvard Apparatus, Holliston, MA) and bacteriostatic 0.9% saline, respectively. Clozapine N-Oxide (CNO) was obtained from Toronto Research Chemicals (North York, ON) and dissolved in 1% DMSO in bacteriostatic 0.9% saline. Fluoro-exendin-4 was purchased from AnaSpec (Fremont, CA) and dissolved in bacteriostatic 0.9% saline. The doses and time courses of administration for each of the aforementioned compounds were based on the following experiments in rats: exendin-(9-39)^14^, exendin-4^8,18,19^, CNO^14,15^, and fluoro-exendin-4^20,21^.

### Catheterization Surgery

Rats were handled daily and allowed one week to acclimate to their home cages upon arrival. Prior to surgery, rats were anesthetized with 100 mg/kg ketamine (Midwest Veterinary Supply, Valley Forge, PA) and 10 mg/kg xylazine (Sigma-Aldrich/RBI, St. Louis, MO). An indwelling catheter (SAI Infusion Technologies, Lake Villa, IL) was inserted into the right jugular vein and sutured in place. The catheter was routed to a mesh backmount that was implanted subcutaneously above the shoulder blades. To prevent infection and maintain patency, catheters were flushed daily with 0.2 ml of the antibiotic Timentin (0.93 mg/ml; Fisher, Pittsburgh, PA) dissolved in heparinized 0.9% saline (Butler Schein, Dublin, OH). When not in use, catheters were sealed with plastic obturators.

### Cocaine self-administration, extinction, and the reinstatement of cocaine-seeking behavior

Rats were allowed 7 days to recover from surgery before behavioral testing commenced. Initially, rats were placed in operant conditioning chambers and allowed to lever-press for intravenous infusions of cocaine (0.245 mg/kg/infusion, infused over 5 s) on a fixed-ratio 1 (FR1) schedule of reinforcement similar to our previous studies^8,14,18,22^. Rats were allowed to self-administer a maximum of 30 injections per two-hour operant session. Once a rat achieved at least 20 infusions of cocaine in a single daily operant session under the FR1 schedule, the subject was switched to a fixed-ratio 5 (FR5) schedule of reinforcement. The maximum number of infusions was again limited to 30 per daily self-administration session under the FR5 schedule. For both FR1 and FR5 schedules, a 20 s time-out period followed each cocaine infusion, during which time active lever responses were tabulated but had no scheduled consequences. Responses made on the inactive lever, which had no scheduled consequences, were also recorded. Following 21 days of daily cocaine self-administration sessions, drug-taking behavior was extinguished by replacing the cocaine solution with 0.9% saline. Daily extinction sessions continued until responding on the active lever was <15% of the total active lever responses completed on the last day of cocaine self-administration. Typically, it took ∼5-7 days for rats to meet this criterion. Once cocaine self-administration was extinguished, rats entered the reinstatement phase of the experiment. To reinstate cocaine-seeking behavior, rats received an acute priming injection of cocaine (10 mg/kg, i.p.) immediately prior to a two-hour reinstatement test session^8,14,18^. During reinstatement test sessions, satisfaction of the response requirement (i.e., five presses on the active lever) resulted in an infusion of saline. Using a between-sessions design, each reinstatement test session was followed by extinction sessions until responding was again <15% of the total active lever responses completed on the last day of cocaine self-administration. Generally, 1-2 days of extinction were necessary to reach extinction criterion between reinstatement test sessions.

### Blood Collection and Analysis

To determine the effects of cocaine taking and subsequent abstinence on circulating GLP-1 levels, a group of Sprague-Dawley rats underwent catheterization surgery and were then randomly assigned to one of two treatment groups: cocaine-experimental group or yoked saline controls. Cocaine-experimental rats were allowed to self-administer cocaine on an FR1 schedule of reinforcement for 21 days. Each rat that was allowed to respond for contingent cocaine infusions was paired with a yoked rat that received infusions of saline. While lever pressing for the yoked saline rats had no scheduled consequences, these rats received the same number and temporal pattern of infusions as self-administered by their paired cocaine-experimental rat. Blood samples were collected from the tail immediately after operant sessions on self-administration day 21, extinction day 1, and extinction day 7. Blood was collected into SAFE-T-FILL capillary collection tubes (RAM Scientific) on ice and then centrifuged at 4°C for 10 min at 14,000 rpm (Centrifuge 5804R, Eppendorf). Plasma was aliquoted into Eppendorf tubes and stored at -80°C. Plasma GLP-1 concentrations were analyzed by an experimenter blinded to treatment conditions in the DRC Radioimmunoassay and Biomarkers Core at Penn using the Human GLP-1(7-36) ELISA Kit (ab184857; Abcam) according to manufacturer’s instructions. Briefly, plasma was diluted 1:10 with Sample Diluent, and 50 μl each of diluted plasma and Antibody Cocktail Solution were added to each well of a microplate. After 1 h incubation at room temperature, microplates were rinsed three times with Wash Buffer, and then allowed to react with 100 μl of Tetramethylbenzidine Development Solution for 15 minutes. Reactions were terminated by adding 100 μl of Stop Solution to each well and fluorescence was measured using a ELx800 NB University Microplate Reader. Each sample was run in triplicate and the assay sensitivity was 25 pg/mL. Mean GLP-1 levels are expressed as picograms per millimeter of plasma.

### Chemogenetic activation of NTS→VTA projections in cocaine-experienced rats

To selectively target NTS→VTA projections, Sprague-Dawley rats underwent catheterization surgery and were immediately mounted in a stereotaxic apparatus (Kopf Instruments, CA). Rats received bilateral infusions of the retrogradely infecting canine adenovirus-2 expressing Cre recombinase (CAV2-Cre) directly into the VTA (-6.00 mm A/P, ± 0.50 mm M/L, and -8.50 mm D/V relative to bregma according to the atlas of Paxinos and Watson^23^). During the same surgical session, an AAV expressing a neural activating DREADD (AAV2-hSyn-DIO-hM3D(Gq)-mCherry) or control virus (AAV2-hSyn-DIO-mCherry) was infused bilaterally into the caudal NTS (-1.00 mm A/P relative to the occipital fissure; ±0.50 mm M/L and -6.00 mm D/V relative to bregma according to the atlas of Paxinos and Watson^23^.) Both viruses were infused at a titer of 1 x 10^12^ gc/ml in a volume of 500 nl over 90 secs. Microinjectors were left in place for an additional 90 sec after infusion to allow for diffusion away from the injection sites. We have used this dual virus approach before to dissect the functional role of specific neural circuits in cocaine seeking^14^. After a 14-day viral expression and recovery period, rats were allowed to self-administer cocaine for 21 days as described above. Once cocaine taking was extinguished, rats were pretreated with vehicle or CNO (0.1 or 1.0 mg/kg, i.p.) 30 min prior to an acute priming injection of cocaine (10 mg/kg, i.p.) similar to our previous reinstatement experiments in rats^14^.

To determine if the effects of activating NTS→VTA projections on cocaine seeking were due to increased GLP-1 signaling in the VTA, a subset of rats were implanted with bilateral VTA guide cannulas (26 gauge; 16 mm; Plastics One, Roanoke, VA) following viral infusions. Guide cannulas were implanted 2.0mm dorsal to the VTA (-6.00 mm A/P, ±0.50 mm M/L, and -6.50 mm D/V relative to bregma according to Paxinos and Western^23^) and cemented in place by affixing dental acrylic to stainless steel screws secured in the skull. An obturator (33 gauge; Plastics One) was inserted into each guide cannula to prevent occlusion. On treatment days, microinjectors that extended 2.0 mm past the ventral ends of the guide cannulas were used to infuse drugs directly into the VTA. Using a within subjects, counterbalanced design, rats were pretreated with bilateral infusions of vehicle or exendin-(9-39) (10 µg/100nl) directly into the VTA 10 min before a systemic injection of vehicle or CNO (1.0 mg/kg, i.p.), similar to our previous experiments^14^. The ability of intra-VTA exendin-(9-39) to block the effects of CNO on cocaine priming-induced reinstatement were assessed 30 minutes later.

Brains were dissected immediately following the last reinstatement test session. Immunohistochemistry was performed on coronal sections of the NTS to 1) visualize viral expression, 2) quantify CNO-induced c-Fos expression in mCherry- and hM3D(Gq)-expressing neurons, and 3) identify DREADD expression in GLP-1-positive NTS cells. Co-localization of mCherry and c-Fos following CNO treatment was quantified using three representative coronal sections of the NTS from each brain (*n*=3 rats/treatment; 3 slices/rat). Missed cannula placements and/or lack of viral expression resulted in rats being excluded from subsequent data analyses.

### Ad libitum food intake

A subset of rats was housed in hanging wire cages with *ad libitum* access to food and water as described previously^8,14,19^. After cocaine reinstatement test sessions, rats were immediately returned to the hanging wire cages and given *ad libitum* access to normal chow (5053 – Pico Lab Rodent Diet 20; LabDiet, Richmond, IN). Food weights were measured 1, 3, 6, and 24 h post session (3, 5, 8, and 26 h post vehicle or CNO injection). Body weight was measured 24 h post session (26 h post CNO injection) similar to our previous studies^14^.

### Verification of cannula placements

After completion of all microinjection experiments, rats were anesthetized with pentobarbital (100 mg/kg, i.p.). Brains were dissected and drop fixed in 10% formalin. Coronal sections (100 µm) were taken at the level of the VTA using a cryostat (Leica 3050S; Leica Corp., Deerfield, IL). An individual blinded to behavioral responses verified microinjection sites using light microscopy. Rats with cannula placements outside of the targeted brain region and/or excessive mechanical damage were excluded from subsequent data analyses.

### Immunohistochemistry

Rats were deeply anesthetized with fatal plus (100 mg/kg, i.p.; Vortech Pharmaceuticals; Dearborn MI) and transcardially perfused with 0.1 M PBS, pH 7.4, followed with 4% formalin in 0.1 M PBS. Brains were removed, postfixed overnight in 4% formalin in 0.1 M PBS, and then cryoprotected in 20% sucrose in 0.1 M PBS at 4°C for three days. Coronal sections (30 µm) were taken using a cryostat (Leica 3050S; Leica Corp., Deerfield, IL). Brain sections were stored in 0.1 M PBS at 4°C until processed. Immunohistochemistry was performed on free-floating coronal sections containing the NTS or VTA according to modified procedures from previously published studies^14,24^. Briefly, sections were washed with 1% sodium borohydride followed by 0.1 M PBS. Sections were then blocked in 0.1 M PBS containing 5% normal donkey serum and 0.2% Triton-X for 1 h at room temperature. Sections were incubated in primary antibodies overnight, and then, following a PBS rinse, incubated in secondary antibodies for 2 h. The primary antibodies used were mouse anti-c-Fos (1:500, sc-271243, Santa Cruz Biotechnology, Inc., Dallas, TX), rabbit anti-GLP-1 (1:1000, T-4363, Peninsula Laboratories International, Inc., San Carlos, CA), and mouse anti-mCherry (1:1000, 632543, Takara, Kyoto, Japan). Secondary antibodies were donkey anti-mouse Alexa Fluor 488 (1:500), donkey anti-rabbit Alexa Fluor 488 (1:500), and donkey anti-mouse Alexa Fluor 594 (1:500) from Jackson ImmunoResearch (West Grove, PA). Sections were then washed and mounted onto glass slides and coverslipped using Vectashield (Vector Laboratories; Burlingame, CA). Sections were visualized with a Leica SP5 X confocal microscope using the 63x oil-immersion objectives along with 488 and 594 nm laser lines. Image z-stacks were captured at the 63x oil-immersion objectives with a step size of 1 μm.

### Electrophysiology validating CNO-induced DREADD activation of NTS cells

Rats expressing AAV2-hSyn-DIO-hM3D(Gq)-mCherry or AAV2-hSyn-DIO-mCherry in the NTS were deeply anesthetized with isoflurane then quickly perfused with ice-cold cutting solution containing (in mM): 92 N-methyl-d-glucamine (NMDG), 2.5 KCl, 1.2 NaH_2_PO_4_, 30 NaHCO_3_, 20 HEPES, 25 glucose, 5 sodium ascorbate, 2 thiourea, 3 sodium pyruvate, 10 MgSO_4_, and 0.5 CaCl_2_, saturated with carbogen (95% O2/5% CO2), pH 7.4 with HCl, osmolarity 305-315 mOsm (Ting et al., 2014). Following decapitation, brains were removed for dissection, and horizontal slices (250 μM thick) of the NTS were obtained using a VT1000S vibratome (Leica, Weltzar, Germany). Slices were made at 4°C then transferred to a holding chamber with the same cutting solution, and incubated at 37°C for 10-12 min. Slices were then moved to a beaker of room temperature holding ACSF containing (in mM): 86 NaCl, 2.5 KCl, 1.2 NaH2PO4, 35 NaHCO3, 20 HEPES, 25 glucose, 5 sodium ascorbate, 2 thiourea, 3 sodium pyruvate, 1 MgCl2, and 2 CaCl2, saturated with carbogen, pH 7.3-7.4, osmolarity 305-315 mOsm. Slices were allowed to recover for at least 45 min before performing recordings.

Slices were transferred to a Nikon Eclipse FN1 upright microscope equipped for Differential Interference Contrast (DIC) infrared optics. The recording chamber was continuously perfused with oxygenated recording ACSF containing (in mM): NaCl 119, KCl 2.5, NaHCO3 26, NaH2PO4 1.2, glucose 12.5, HEPES 5, MgSO4 1, CaCl2 2, pH 7.3-7.4, osmolarity 305-315 mOsm. The solution was heated to 32 ± 1°C using an automatic temperature controller (Warner Instruments). The NTS was identified using a 5X objective and individual neurons were magnified with a 40x water immersion lens. DREADD-positive neurons were further identified by fluorescence with a DS-Red filter (Nikon). Recording pipettes were pulled from borosilicate glass capillaries (World Precision Instruments) to a resistance of 3.8-5.0 MΩ when filled with intracellular solution. The intracellular solution contained the following (in mM) potassium gluconate 145, KCl 2.5, NaCl 2.5, BAPTA 0.1, HEPES 10, L-glutathione 1.0, sodium phosphocreatine 7.5, Mg-ATP 2.0, and Tris-GTP 0.25, pH 7.2-7.3 with KOH, osmolarity 285-295 mOsm. All recordings were performed in whole-cell current-clamp mode using a MultiClamp 700B amplifier. Baseline spontaneous firing was recorded for ≥ 5 min prior to the activation of DREADD-positive neurons by CNO (30 μM) dissolved in the recording ACSF. After ≥ 5 min CNO was washed out with regular recording ACSF. All recordings were low-pass filtered at 3kHz, amplified 5 times, and then digitized at 20kHz using a Digidata 1440A acquisition board and pClamp10 software (both from Molecular Devices). For all experiments, access resistance was 15–25 MΩ, uncompensated, and monitored continuously during recording. Cells with a change in access resistance >20% over the course of data acquisition were not accepted for data analysis.

### Fluorescence in situ hybridization (FISH)

FISH was used to quantify cell type-specific *Glp1r* expression in the VTA of drug-naïve, wild type Long Evans rats (*n*=3 rats) and verify cell-type specific GCaMP8f expression in transgenic rats used in the fiber photometry experiments (*n=*4 rats; 2 rats/transgenic line). Brains were dissected, flash frozen in -20°C isopentane, and stored at -80°C. Using a cryostat (Leica 3050S; Leica Corp., Deerfield, IL), coronal sections (8 µm) were taken at the level of the VTA and immediately mounted onto Superfrost Plus slides (Fisher Scientific). *Glp1r*, *Th*, and *Gad1* mRNA transcripts were detected using the RNAScope Multiplex Fluorescent Reagent Kit V2 (Cat. 320850; Advanced Cell Diagnostics (ACD), Hayward, CA) per manufacturer’s protocol. Slide mounted sections were rinsed in 10% formaldehyde for 15 min at 4°C. Following dehydration in ascending concentrations of ethanol solutions (5 min washes in 50%, 70%, 100%, and 100% ethanol), slides were air-dried, and a hydrophobic barrier was created around the sections using a hydrophobic pen (Vector Labs). The sections were then rinsed with PBS and treated with Protease IV (Advanced Cell Diagnostics (ACD), Hayward, CA) for 30 min at RT (followed by two 1 min rinses in PBS).

Pretreated tissue sections were processed immediately using probes designed by ACD to detect *Glp1r* (Rn-Glp1r-C1; 315221) or e*Gfp* (EGFP; 400281-C1), *Gad1* (Rn-Gad1-C2; 316401-C2), and *Th* (Rn-Th-C3; 314651-C3) transcripts. Sections were incubated in a cocktail containing probes (Probe dilution: 50C1:1C2:1C3) for 2 h at 40°C in a HybEZTM oven (ACD). Following signal amplification, opal dyes [Opal 520 (OP-001001), Opal 570 (OP-001003), and Opal 690 (OP-001006) (Akoya, Marlborough, MA)] were diluted in TSA buffer (1:1000) and applied to the sections. After the final wash, slides were coverslipped using Fluorogel mounting medium with DAPI (Fisher Scientific). Brain sections at four levels relative to bregma (-5.30, -5.60, -6.04, and -6.30mm A/P) were sampled from each rat. A total of four images (one image per level) from each rat were captured with a Keyence fluorescence microscope using the 10x, 20x, and 60x oil-immersion objectives. Image z-stacks were collected with a step size of 2 µm. To quantify *Glp1r* expression in *Gad1-* and *Th*-expressing cells, the number of cells co-labeled for *Glp1r* and *Gad1* transcripts or *Th* transcripts in 10x images were counted at each level per rat and totaled (*n=*3 rats; 4 slices/rat). These numbers were divided by the total number of cells that expressed *Glp1r* to calculate the percentage of *Glp1r-* expressing cells that co-express *Gad1* and/or *Th* transcripts at different A/P levels and in total. The number of cells expressing *Gad1* or *Th* transcripts were also counted at each level to quantify the total number of *Th* and *Gad*-expressing cells in the VTA.

### Single nuclei RNA-sequencing

Adult male Sprague-Dawley rats (*n*=5) were rapidly decapitated, their brains removed, and flash frozen in dry ice cold isopentane. Bilateral VTA punches were prepared on a cryostat and combined into a single sample. Nuclei were isolated from the samples as described previously^25,26^. Nuclei were processed for the 10x Genomics 3’ gene expression assay (v3.1) and sequenced per manufacturer protocols. Raw sequencing data was processed and aligned to the rat reference genome as described previously^27^. Downstream analyses of count data were performed with Seurat v4.3.0. Nuclei with mitochondrial transcript fractions ≥5% or with ≤200 genes detected were removed and the remaining nuclei were clustered at a resolution of 0.1. Count matrices were then corrected for cell-free mRNA background noise using SoupX v1.6.2 and nuclei with ≥5,000 genes detected in the corrected data were removed. Count data was then normalized with SCTransform and all samples were integrated with Seurat. Doublets were identified with scDblFinder v1.12.0 and removed. Clusters with low UMI or displaying markers for more than one major cell type were also removed. The final dataset was clustered at a resolution of 0.08 using the first 10 principal components based on highly variably expressed genes and clusters were assigned to major cell types based on known marker genes.

### Dual-Wavelength in vivo Fiber Photometry Recordings

465nm and 405nm LEDs (Tucker Davis Technologies (TDT), Alachua, FL) were used simultaneously in each rat to excite GCaMP8f and measure intracellular Ca^2+^ dynamics. The 465nm LED excites (signal) calcium-dependent GCaMP8f fluorescence to provide a proxy for neural activity, while the 405nm LED (isosbestic) excites calcium-independent GCaMP8f fluorescence to control for movement and fiber bleaching. The intensities of the 465nm and 405nm LEDs were sinusoidally modulated at 210Hz and 330Hz, respectively. Both LED light sources were coupled to a dichroic mirror containing filter cube (FMC4, Doric Lenses, Quebec, Canada) and converged onto a fiber optical patch cord (Doric Lenses) attached to a ferule-capped optic implant (Doric Lenses). The emitted 510nm fluorescent signal was collected through the same fiber optic patch cord coupled to the LEDs and filtered before being collected and amplified by a lock-in amplifier at an acquisition rate of 6 Hz (RZ10x, TDT). Using the iCON control interface and Synapse recording software (TDT), the RZ processor was integrated with operant chambers to record from freely moving rats during cocaine reinstatement test sessions. A custom closed-loop experiment was designed in Pynapse (TDT), a Python-based programming environment, to conduct drug seeking experiments, record data, and time stamp behavioral events (e.g., lever presses).

### Measuring cell type-specific Ca^2+^ dynamics in VTA neurons during cocaine reinstatement

To identify the midbrain mechanisms underlying the efficacy of exendin-4 on cocaine seeking, transgenic rats expressing Cre recombinase under the *Gad1* or *Th* promoter were implanted with indwelling jugular catheters as described above. A Cre-dependent GCaMP8f virus (AAV9-Syn-Flex-GCaMP8f-WPRE-SV40) was infused directly into the VTA (*Th* rats: -6.00 mm A/P, +or-0.50 mm M/L, and -8.50 mm D/V; *Gad1* rats: -6.30 mm A/P, +or- 0.50 mm M/L, -8.50 mm D/V). These coordinates were based on subregion- and cell type-specific patterns of *Gad1* and *Th* expression in the VTA (Suppl. Fig 1A & 1B). Viruses were infused unilaterally in a volume of 1μL over 5 minutes (titer = 1.0 x10^12^ gc/ml). Microinjectors were left in place for an additional 5 minutes after infusion to allow for diffusion away from the injection sites. A ferrule-capped optic fiber (Doric Lenses) was implanted 10 μm above the viral injection site (*Th* rats: -6.00 mm A/P, +or- 0.50 mm M/L, and - 8.49 mm D/V; *Gad1* rats: -6.30 mm A/P, +or- 0.50 mm M/L, and -8.49 mm D/V) and secured to the skull with Metabond cement (Parkell) and dental acrylic.

After a 14-day viral expression and recovery period, rats were allowed to self-administer cocaine as described above. Drug taking was then extinguished by replacing cocaine with saline. Once cocaine taking was extinguished, rats were pretreated with vehicle or exendin-4 (2.0 μg/kg, i.p.) 10 minutes prior to a priming injection of cocaine (10 mg/kg, i.p.). The dose of exendin-4 tested was shown previously to reduce drug taking in cocaine-experienced rats^8^. Following the acute priming injection of cocaine, a fiber optic patch cord was secured to the optic fiber-containing ferrule implant. Rats were then placed immediately into the operant chambers and drug seeking was assessed during a two-hour reinstatement test session. Calcium-dependent GCaMP8f fluorescence was recorded for the first 30 minutes of the two-hour reinstatement test session and lever-dependent behavioral events were timestamped using the iCON control interface and Pynapse (TDT). Signals associated with the first three response requirements completed (i.e., the fifth, tenth, and fifteenth active lever presses) were analyzed in this study. Signals associated with inactive lever presses were also recorded and analyzed. Cell type-specific Ca^2+^ responses were also measured in separate rats that self-administered saline to determine if the observed effects on calcium dynamics were specific to cocaine-seeking behavior and not due to handling and/or exposure to the operant chambers.

### Fiber Photometry Data Analysis

Raw signal (465 nm) and isosbestic control (405 nm) channel data were exported from Synapse to a CSV file using a MATLAB script provided by TDT. Custom R scripts were then used to independently process each trace (https://github.com/rileymerkel/Schmidt-Lab.git). First, the scale of channels was normalized using a least-squares regression to find the relationship between the signal and control channel. The returning slope and intercept were used to generate a scaled control channel. ΔF/F was then generated by subtracting the fitted isosbestic control channel from the signal channel to eliminate any movement or bleaching artifacts. To identify changes in fluorescence associated with cocaine seeking, time stamps were used to divide the processed trace into 10 second trials. Trials were z-scored to a baseline period defined as 5 seconds prior to engagement with the operant box levers.

### Statistics

For all cocaine reinstatement experiments, total active and inactive lever responses were analyzed with repeated measures (RM) two-way mixed-factors ANOVAs. Cumulative food intake data were also analyzed with RM two-way mixed-factors ANOVAs. For fiber photometry experiments, area under the curve (AUC) and maximum z-scores were analyzed with separate RM two-way mixed-factors ANOVAs. Pairwise analyses were made using Bonferroni *post-hoc* tests (*p*<0.05). Findings from all other experiments were analyzed using two-tailed, paired or unpaired *t*-tests (*p*<0.05).

### Data availability

All relevant data are included in the paper and its supplementary figures. All data generated in this study are provided in the Source Data File. snRNA-seq count matrices for our discovery dataset are available at the NCBI Gene Expression Omnibus (GEO) under accession number GSE######.

### Code availability

The custom code used to analyze the fiber photometry data in this paper is available at (https://github.com/rileymerkel/Schmidt-Lab.git).

## Results

### Voluntary cocaine taking decreases plasma GLP-1 levels

To determine if voluntary cocaine taking and subsequent abstinence alter plasma GLP-1 levels, blood samples were drawn from cocaine-experienced rats and their corresponding yoked-saline controls at three time points (Fig 1A). Plasma GLP-1 concentrations (pg/mL) were significantly decreased in cocaine-experienced rats versus yoked-saline controls following 21 days of self-administration (Fig 1B) and acute withdrawal (i.e., extinction day 1) (Fig 1C). In contrast, there were no significant differences in plasma GLP-1 levels between cocaine-experienced rats and yoked-saline controls after prolonged withdrawal (i.e., extinction day 7) (Fig 1D). These findings, together with our previous studies showing that systemic and intra-cranial infusions of a GLP-1R agonist attenuated cocaine reinstatement^8,14,18^, support the hypothesis that decreased endogenous GLP-1 signaling facilitates/promotes cocaine-taking and -seeking behaviors^5^.

**Figure 1.**
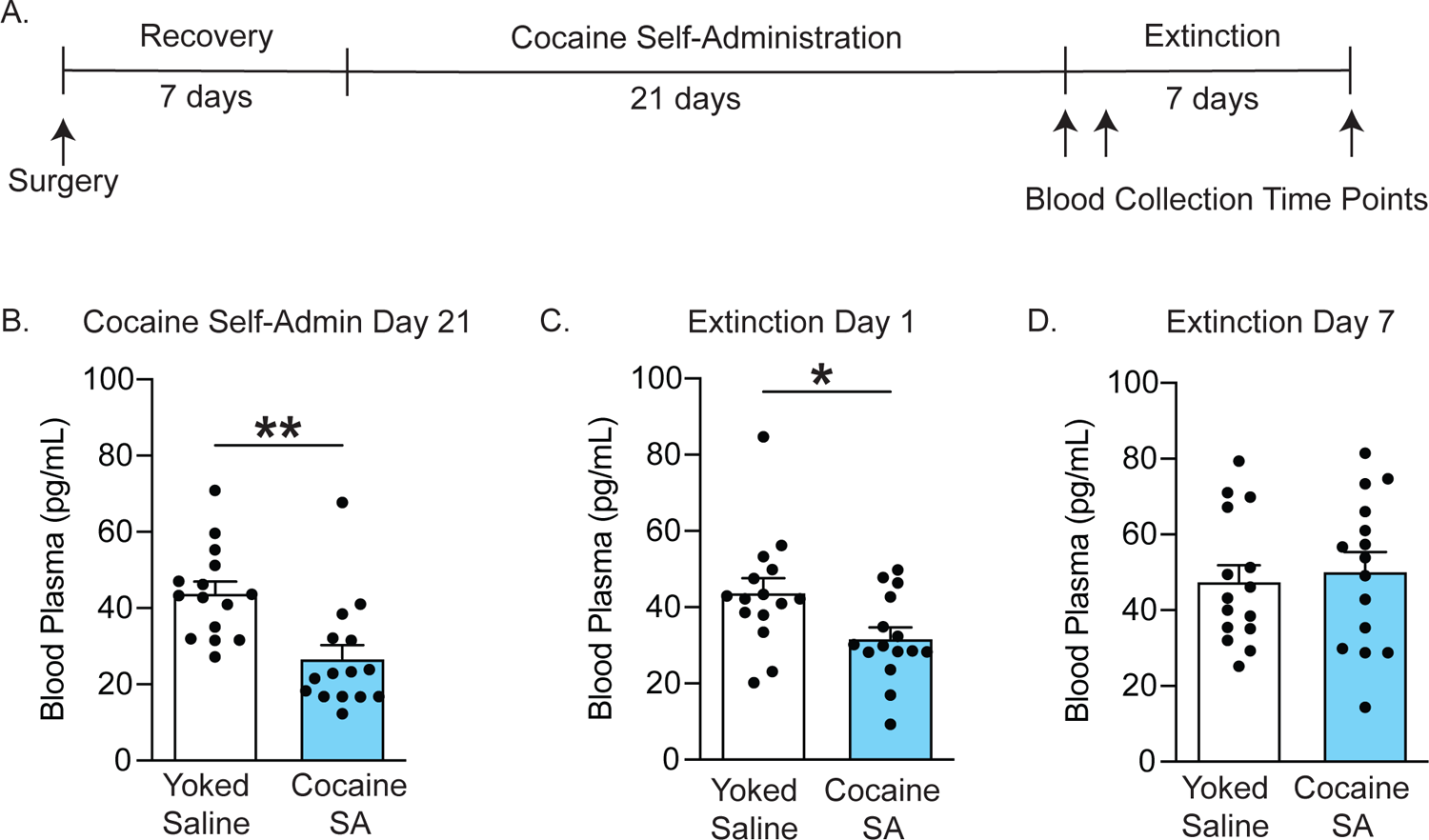
Cocaine self-administration decreases circulating GLP-1 levels. A Schematic illustrating the experimental timeline and blood collection time points (yoked saline: *n* = 15; cocaine self-administration: *n* = 15). **B, C** Plasma GLP-1 concentrations were significantly decreased in cocaine-experienced rats following 21 days of cocaine self-administration (unpaired *t-*test: t_28_ = 3.604, \*\**p* = 0.0012) and 1 day of extinction (unpaired *t-*test: t_28_ = 2.474, *\*p* = 0.0197). **D** There were no significant differences in plasma GLP-1 levels between cocaine-experienced rats and yoked-saline controls following seven days of extinction (unpaired *t*-test: t_28_ = 0.4042, *p* = 0.6891). Data are mean ± SEM.

### Activation of NTS neurons that project to the VTA selectively attenuates drug seeking in cocaine-experienced rats

Despite growing evidence that GLP-1R agonist pharmacotherapy is sufficient to reduce cocaine seeking^8,18^, no studies have investigated the role of endogenous GLP-1-producing neural circuits in drug-seeking behavior. To selectively activate endogenous GLP-1-producing NTS neurons that project to the VTA, the retrogradely infecting virus CAV2 expressing Cre recombinase (CAV2-Cre) was infused into the VTA and a Cre-dependent virus expressing a neural activating DREADD (AAV-DIO-hM3D(Gq)-mCherry) was infused into the NTS (Fig 2A & 2B). IHC was performed on NTS sections to label c-Fos after CNO injection in both hM3D(Gq)-expressing rats and mCherry control rats (Fig 2C). CNO significantly increased c-Fos expression in ∼55% of NTS hM3D(Gq)-expressing neurons compared to <5% of mCherry-expressing neurons (Fig 2D). Whole-cell current clamp recordings were conducted to further validate the ability of CNO to increase activity of hM3D(Gq)-expressing NTS neurons compared to mCherry-expressing control neurons. CNO increased NTS cell firing in hM3D(Gq)-expressing rats (Fig 2E) and had no effect on NTS cell firing in mCherry control rats (Fig 2F).

**Figure 2.**
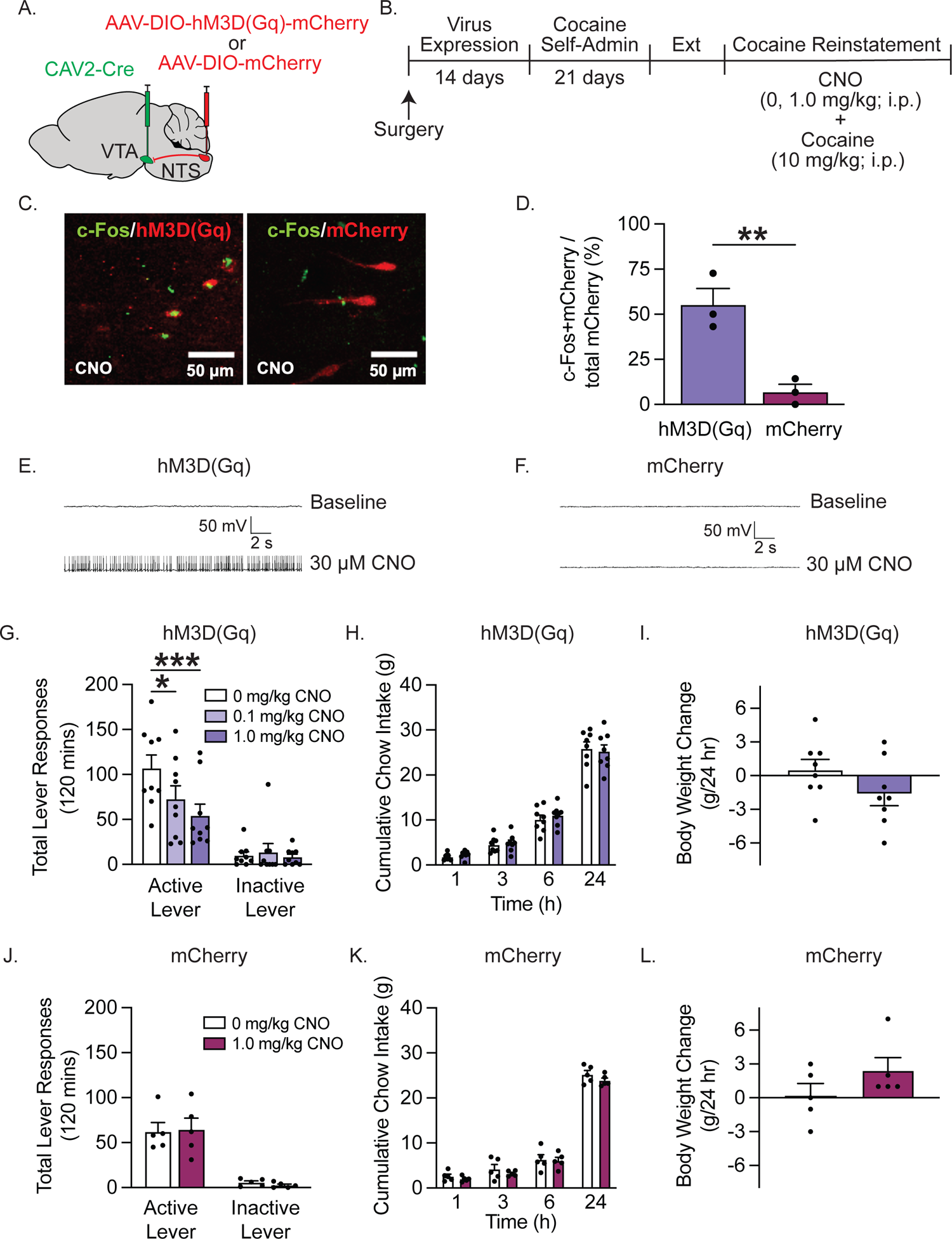
Activating NTS→VTA projections attenuates cocaine seeking. **A, B** Illustration of viral approach and experimental timeline wherein CAV2-Cre and a Cre-dependent AAV expressing hM3D(Gq) or mCherry were infused into the VTA and NTS, respectively, prior to the cocaine self-administration phase of the experiment. **C** Representative images showing increased c-Fos expression in hM3D(Gq)-expressing NTS neurons of rats treated with CNO versus mCherry controls. **D** CNO significantly increased c-Fos expression in ∼55% of hM3D(Gq)-expressing NTS neurons versus <5% of mCherry-expressing NTS neurons *(n* = 3/treatment; unpaired *t*-test: t_4_ = 4.911, \*\**p* = 0.0080). **E, F** Representative whole-cell current-clamp traces showing increased firing of hM3D(Gq)-expressing NTS neurons and no change in firing of mCherry-expressing NTS neurons following CNO administration. **G** CNO dose-dependently decreased active lever presses in rats expressing hM3D(Gq) in NTS neurons that project to the VTA (*n* = 9; two-way RM ANOVA, treatment x lever interaction: F_2_,_16_ = 5.362, *p* = 0.0165; Bonferroni’s test: vehicle vs 0.1 mg/kg CNO, \**p* = 0.025; vehicle vs 1.0 mg/kg CNO, \*\*\**p* = 0.0009). **H, I** CNO did not alter cumulative chow intake (two-way RM ANOVA, treatment: F_1,7_ = 0.1214, *p* = 0.7377; time: F_3,21_ = 184.6, *p* < 0.0001; time x treatment: F_3,21_ = 0.6343, *p* = 0.6012) or 24 hour body weight gain (paired *t*-test: t_7_ = 1.838, *p* = 0.1087) in rats expressing hM3D(Gq) in NTS neurons that project to the VTA (*n =* 8/treatment). **J, K, L** CNO had no effects on cocaine seeking (two-way RM ANOVA, treatment: F_1,4_ = 0.01472, *p* = 0.9093; lever: F_1,4_ = 30.56, *p* = 0.0052; treatment x lever: F_1_,_4_ = 0.6170, *p* = 0.4761), cumulative chow intake (two-way RM ANOVA, treatment: F_1,4_ = 1.435, *p* = 0.2970; time: F_3,12_ = 544.7, *p* < 0.0001; treatment x lever: F_3_,_12_ = 0.421, *p* = 0.7419), or 24 hour body weight gain (paired *t*-test: t_4_ = 2.400, *p* = 0.0743) in mCherry-expressing control rats (*n* = 5). Data are mean ± SEM.

To determine the functional role of NTS→VTA projections in cocaine seeking, rats were pretreated with vehicle or CNO (0.1 or 1.0 mg/kg, i.p.) prior to a cocaine priming-induced reinstatement test session. CNO dose-dependently attenuated cocaine seeking in hM3D(Gq)-expressing rats (Fig 2G). A previous study showed that activation of GLP-1-producing NTS neurons that project to the VTA transiently reduced food intake in drug-naïve mice^28^. In addition, GLP-1R agonists are known to decrease food intake and produce nausea/emesis in both drug-naïve rodents and humans^4,20,29,30^. We screened for these potential adverse effects in our cocaine-experienced rats and found no effects of CNO on cumulative chow intake (Fig 2H) and 24 h body weight gain (Fig 2I) on reinstatement test days. Together, these studies indicate that activation of NTS→VTA projections is sufficient to reduce cocaine seeking during abstinence and that drug seeking is more sensitive to central GLP-1 modulation than non-drug motivated behaviors in cocaine-experienced rats.

Although CNO is often used as the inert ligand for DREADDs, studies show minimal, yet significant, reverse metabolism to clozapine, an active metabolite which could bind to non-DREADD receptors and produce off-target effects on behavior^31^. To control for this, a separate group of rats was infused with the control virus AAV-DIO-mCherry into the NTS and allowed to self-administer cocaine. In contrast to the effects seen in hM3D(Gq)-expressing rats, CNO had no effect on cocaine seeking in rats infused with the mCherry control virus (Fig 2J). These findings indicate that CNO itself does not alter drug-seeking behavior during abstinence following cocaine self-administration. Moreover, there were no effects of CNO on cumulative chow intake (Fig 2K) and 24 h body weight gain (Fig 2L) in mCherry control rats. Thus, the suppressive effects of activating NTS→VTA projections on cocaine seeking are not due to off-target effects of CNO or a CNO metabolite.

### Activation of NTS→VTA projections attenuates cocaine seeking via a GLP-1-dependent mechanism of action in the VTA

Our next goal was to determine if the suppressive effects of activating NTS→VTA projections on cocaine seeking were due to increased GLP-1 signaling in the VTA. Rats were infused with CAV2-Cre into the VTA, AAV-DIO-hM3D(Gq)-mCherry into the NTS, and implanted with guide cannula aimed at the VTA (Fig 3A). Rats then underwent cocaine self-administration, extinction, and reinstatement (Fig 3B). IHC analyses revealed hM3D(Gq) colocalized with GLP-1 in NTS neurons that project to the VTA (Fig 3C). Prior to reinstatement test sessions, the GLP-1R antagonist exendin-(9-39) (10 µg/100nl) was infused directly into the VTA 10 minutes before administration of CNO (1.0 mg/kg, i.p.) to determine if pharmacological inhibition of VTA GLP-1Rs is sufficient to block the suppressive effects of chemogenetic activation of NTS→VTA projections on cocaine seeking (Fig 3D). Intra-VTA exendin-(9-39) pretreatment blocked the ability of CNO to attenuate cocaine seeking (Fig 3E & 3F) indicating that the suppressive effects of activating endogenous NTS→VTA circuits on drug seeking depend, in part, on enhanced GLP-1 signaling in the midbrain.

**Figure 3.**
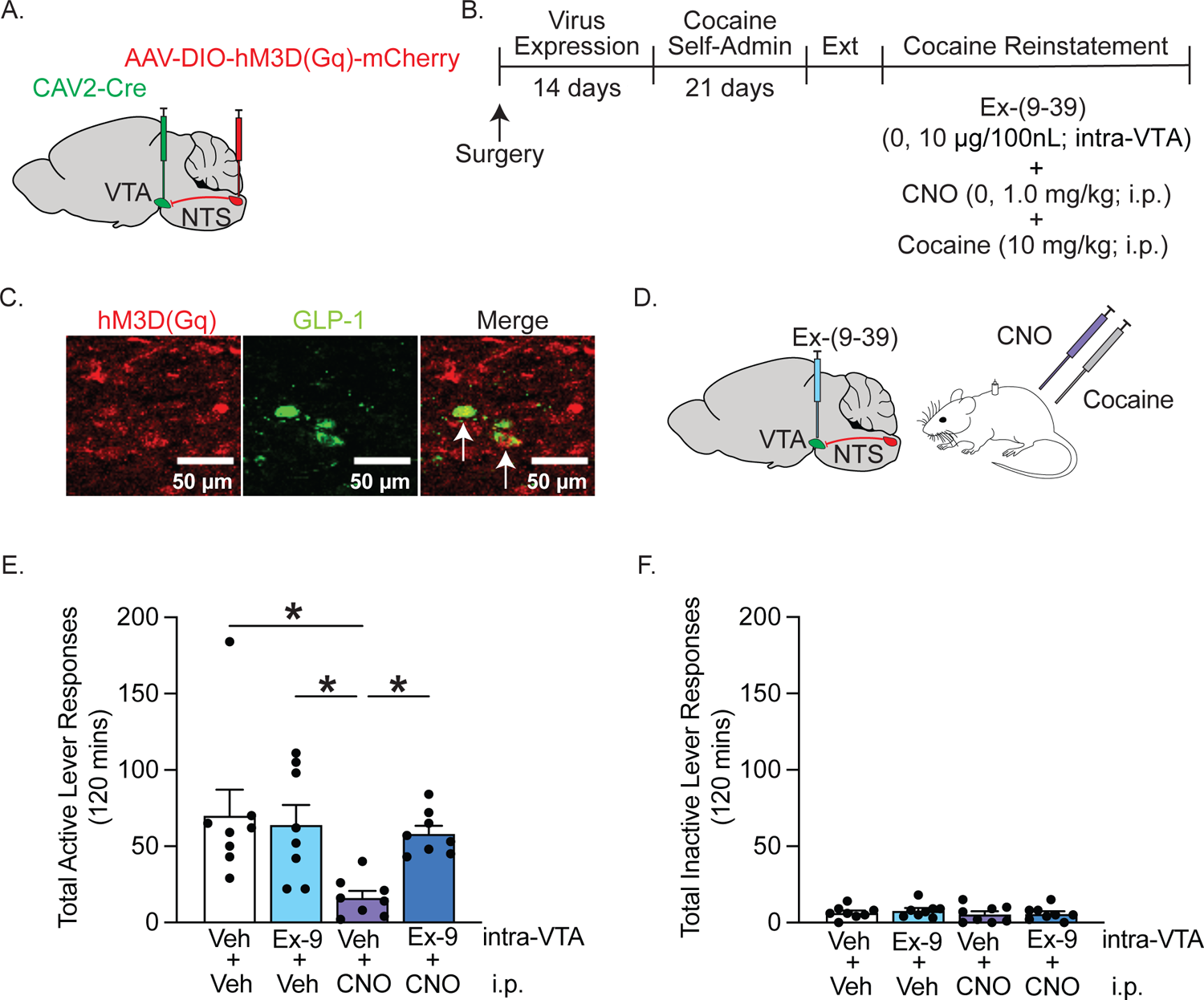
Pharmacological inhibition of VTA GLP-1Rs blocks the suppressive effects of activating NTS→VTA projections on cocaine seeking. **A, B** Illustration of viral approach and experimental timeline wherein CAV2-Cre and a Cre-dependent AAV expressing hM3D(Gq) were infused into the VTA and NTS, respectively, prior to the cocaine self-administration phase of the experiment. **C** Representative images of hM3D(Gq)-expressing NTS neurons that co-express GLP-1. **D** Schematic depicting reinstatement test session treatment conditions. Exendin-(9-39) was infused into the VTA prior to activating endogenous NTS→VTA projections with CNO. **E** Intra-VTA exendin-(9-39) prevented the ability of CNO to suppress active lever presses during reinstatement test sessions *(n* = 8; two-way RM ANOVA, systemic treatment x intra-VTA treatment: F_1_,_7_ = 8.775, *p* = 0.0210; Bonferroni’s test: Veh/Veh vs Veh/CNO, \**p* = 0.0132; Veh/CNO vs Ex-(9-39)/Veh, \**p* = 0.0249; Veh/CNO vs Ex-(9-39)/CNO, \**p* = 0.0441). **F** Intra-VTA exendin-(9-39) had no effect on inactive lever responses (*n* = 8; two-way RM ANOVA, systemic treatment: F_1_,_7_ = 2.333, *p* = 0.1705; intra-VTA treatment: F_1,7_ = 0.320, *p* = 0.5891; systemic treatment x intra-VTA treatment: F_1_,_7_ = 0.138, *p* = 0.7207). Data are mean ± SEM.

### GLP-1Rs are expressed primarily on GABAergic cells in the VTA

Next, we showed that a behaviorally relevant dose of exendin-4 that reduced cocaine seeking^8,18^ crossed the blood brain barrier, bound putative GLP-1Rs on neurons in the VTA, and induced c-Fos expression in these neurons (Suppl. 2A). The exact cell types expressing GLP-1Rs in the VTA, however, remain unclear. To further investigate the mechanisms by which GLP-1R agonist pharmacotherapy alters midbrain neuron activity and attenuates cocaine seeking, we used FISH to phenotype GLP-1R-expressing cells in the VTA. Interestingly, we found that ∼90% of *Glp1r-*expressing cells co-express *Gad1*, and that no *Glp1r* transcripts were detected in *Th-*positive cells (Fig 4A & 4B). In addition, ∼10% of *Glp1r*-expressing cells belonged to non-identified cell type(s) (Fig 4B). When quantifying expression throughout the rostral-caudal axis of the VTA, FISH analyses revealed a greater number *Glp1r*-expressing, *Gad1*-positive cells in the posterior VTA when compared to more anterior subregions of the VTA (Fig 4C & 4D). To validate the expression pattern of *Glp1r*, we performed snRNA-seq on rat VTA samples. Neurons and all major glial populations were identified in the

**Figure 4.**
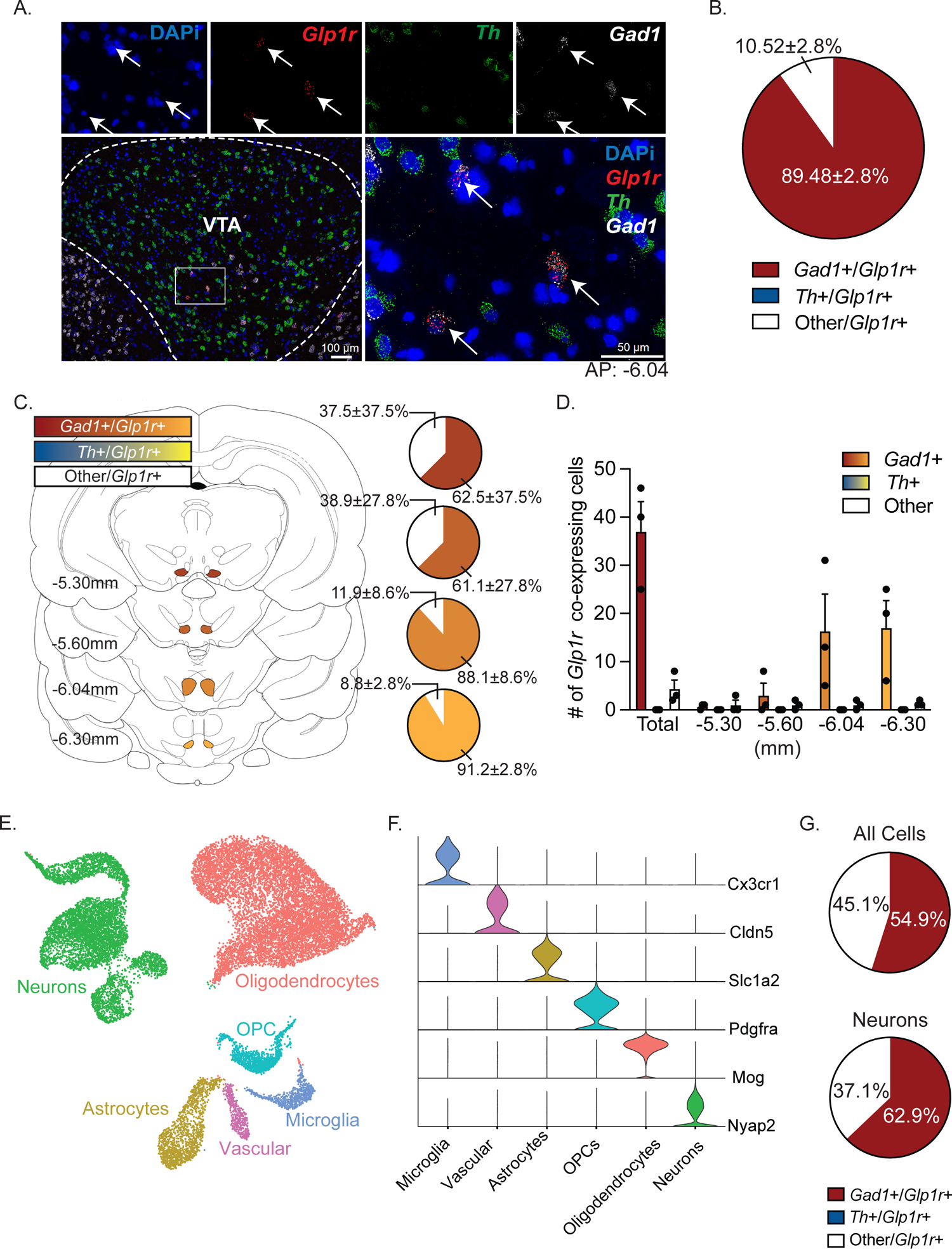
GLP-1Rs are expressed primarily on GABAergic neurons in the VTA. A FISH revealed *Glp1r* transcripts co-expressed with *Gad1* transcripts in the VTA [DAPi (blue), *Glp1r* (red), *Th* (green), and *Gad1* (white)]. **B** ∼90% of all *Glp1r*-expressing neurons in the VTA co-express *Gad1.* No *Glp1r* transcripts were detected in *Th*-positive cells (*n =* 3 rats; 4 slices per rat). **C, D** Both the total percentage and number of *Glp1r*-positive cells co-expressing *Gad1* are greater in posterior regions of the VTA (*n* = 3 rats). **E** Uniform manifold approximation and projection (UMAP) dimension reduction plot for snRNA-seq from the VTA of drug-naïve rats (*n* = 5). Nuclei are colored by major cell type. **F** Violin plot showing normalized expression of marker genes for major VTA cell types. **G** Pie charts displaying the percentage of *Glp1r*-expressing cells (top) or neurons (bottom) that co-express *Gad1*, *Th*, or neither marker in the snRNA-seq dataset. Data are mean ± SEM.

snRNA-seq dataset (Fig 4E & 4F). Consistent with our FISH results, the majority of *Glp1r*-expressing nuclei also expressed *Gad1* and there was no observed overlap between *Glp1r* and *Th* expression (Fig 4G). These anatomical studies clearly showed that GLP-1Rs are expressed primarily on GABA neurons in the VTA and support the hypothesis that GLP-1R agonists increase inhibitory GABA transmission in the VTA to reduce cocaine seeking.

### Systemic GLP-1R agonist pharmacotherapy increases activity of VTA GABA neurons and attenuates cocaine seeking

Our FISH and snRNA-seq studies revealed GLP-1Rs expressed primarily on GABA neurons in the VTA (Fig 4). However, it is unclear how GLP-1R activation influences the activity of VTA GABA neurons in cocaine-experienced rats and how these cell type-specific cellular responses are related to cocaine-seeking behavior. We used *in vivo* fiber photometry in transgenic rats to determine how GLP-1R agonist pharmacotherapy alters activity of VTA GABA neurons during reinstatement test sessions. To measure changes in intra-cellular calcium levels in midbrain GABA neurons during cocaine seeking, a Cre-dependent GCaMP8f virus was infused into the VTA of rats expressing Cre-recombinase under the *GAD* promoter and a fiberoptic cannula was implanted directly above the infusion site (Fig 5A). A representative image of subregion-specific viral expression is shown in Figure 5B. Fluorescent labelling of *eGfp, Th,* and *Gad1* mRNA transcripts using FISH confirmed selective GCaMP8f expression in *Gad1*-positive neurons in the VTA (Fig 5B).

**Figure 5.**
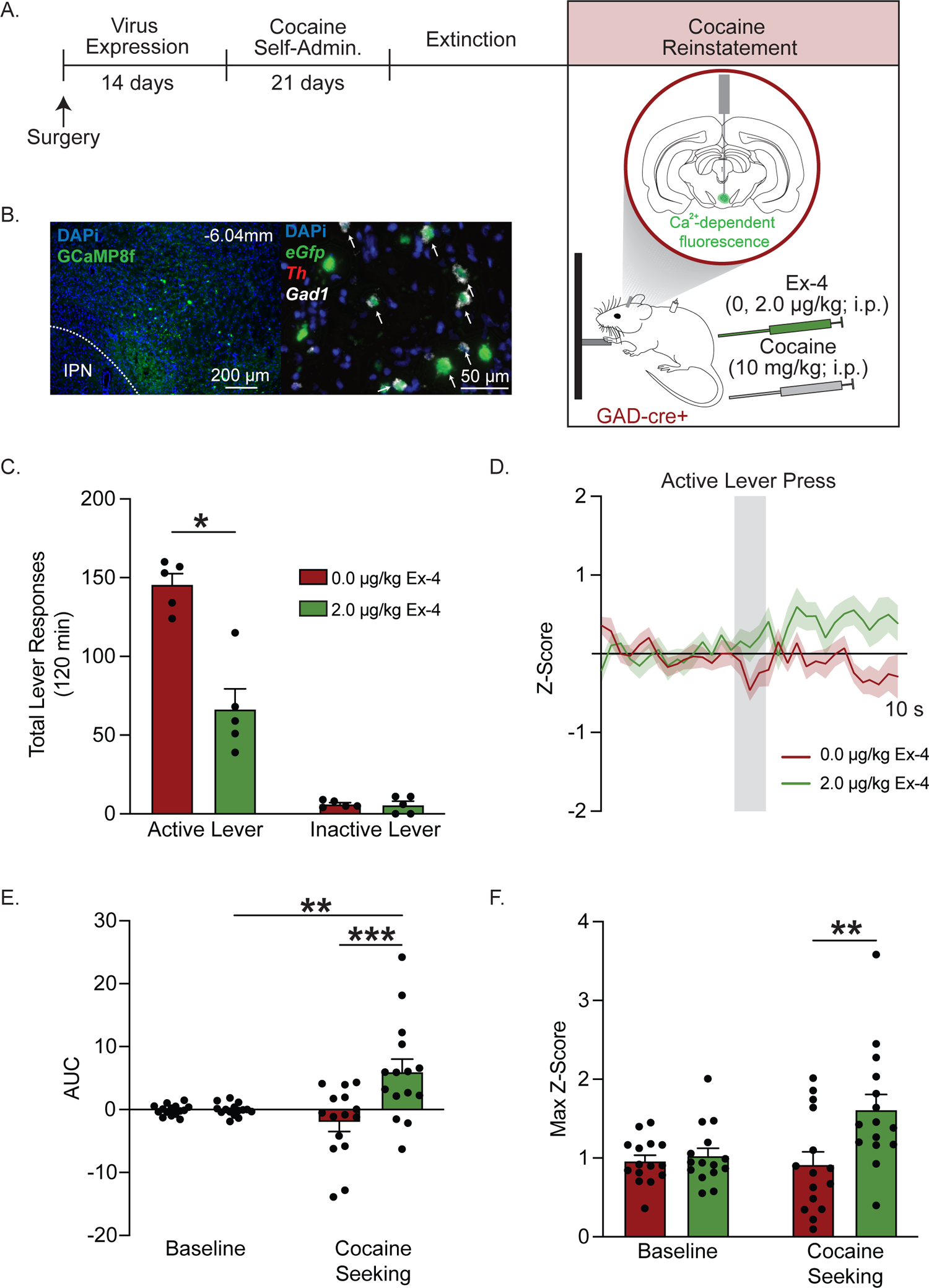
Systemic GLP-1R agonist pharmacotherapy increases VTA GABA neuron activity and decreases cocaine seeking. A Schematic illustrating the viral approach and experimental timeline wherein a Cre-dependent AAV expressing GCaMP8f virus was infused into the VTA of GAD-Cre rats to measure intra-cellular calcium dynamics in VTA GABA neurons during reinstatement tests sessions. **B** Representative image of GCaMP8f viral expression in the VTA. FISH confirmed selective GCaMP8f expression in *Gad1*-positive cells [DAPi (blue), *eGfp* (green), *Th* (red), and *Gad1* (white)]. **C** Systemic exendin-4 administration decreased active lever responses during cocaine reinstatement test sessions (*n =* 5; two-way RM ANOVA, treatment x lever: F_1_,_4_ = 19.64, *p* = 0.0114; Bonferroni’s test: vehicle vs exendin-4, \**p* = 0.0129). **D** Normalized z-score traces during cocaine seeking in rats pretreated with vehicle or exendin-4 *(n* = 5: 3 active lever presses/rat/treatment). **E** Cocaine seeking significantly increased the AUC of recorded Ca^2+^ signals from VTA GABA neurons in rats treated with exendin-4, but not vehicle (*n* = 15; two-way RM ANOVA, treatment x time: F_1_,_28_ = 9.652, *p* = 0.0043; Bonferroni’s test: baseline vs cocaine seeking in rats treated with exendin-4, \*\**p* = 0.0099). The AUC from rats treated with exendin-4 was also significantly increased during cocaine seeking compared to vehicle-treated controls (Bonferroni’s test: 0.0 vs 2.0 μg/kg exendin-4 during cocaine seeking, *** *p* = 0.0001). **F** Exendin-4 significantly increased maximum z-scores in VTA GABA neurons during cocaine seeking compared to vehicle-treated controls (*n* = 15; two-way RM ANOVA, treatment x time: F_1_,_28_ = 4.110, *p* = 0.0522; Bonferroni’s test: 0.0 vs 2.0 μg/kg exendin-4 during cocaine seeking, ** *p* = 0.0036). Data are mean ± SEM.

To determine how GLP-1R activation alters calcium dynamics in VTA GABA neurons during cocaine seeking, rats were pretreated with vehicle or exendin-4 (2.0 µg/kg, i.p.) prior to a cocaine priming-induced reinstatement test session. Consistent with our prior studies^8,18^, exendin-4 significantly reduced cocaine seeking in GAD-Cre rats (Fig 5C). Cell type-specific normalized z-score traces during cocaine seeking are presented in Figure 5D. Area under the curve (AUC) and maximum z-scores were calculated from these traces. AUC was significantly increased during cocaine seeking compared to baseline in VTA GABA neurons of rats pretreated with exendin-4, indicating that cocaine-seeking behavior evokes transient increases in VTA GABA neuron activity in exendin-4-treated rats (Fig 5E & 5F). Importantly, AUC and maximum z-scores were significantly increased in VTA GABA neurons from exendin-4-treated rats compared to VTA GABA neurons from vehicle-treated controls (Fig 5E & 5F). These results indicate that exendin-4 pretreatment produces greater activity of VTA GABA neurons during drug seeking compared to vehicle-treated controls. Normalized z-score traces associated with inactive lever presses during reinstatement test sessions are presented in Supplementary Figure 3A. There were no effects of treatment on AUC and maximum z-scores from VTA GABA neurons following inactive lever responses during reinstatement test sessions (Suppl. Fig 3B & 3C). To confirm that these calcium dynamics were specific to cocaine-seeking behavior, a separate group of GAD-Cre rats was allowed to self-administer saline before undergoing extinction and subsequent reinstatement tests. There were no effects of vehicle or exendin-4 on AUC or maximum z-scores from VTA GABA neurons following active lever presses during the reinstatement test sessions in drug-naïve rats (Suppl. 4A-D).

Together, these findings indicate that GLP-1R agonist pharmacotherapy increases calcium dynamics in VTA GABA neurons during cocaine seeking, effects associated with decreased drug-seeking behavior during withdrawal.

### Systemic GLP-1R agonist pharmacotherapy reduces activity of VTA dopamine neurons and attenuates cocaine seeking

Systemic GLP-1R agonist administration reduced cocaine-evoked dopamine release in the nucleus accumbens^32^, a downstream target of VTA dopamine neurons known to play a critical role in cocaine seeking^33,34^. Given that GLP-1R agonist pharmacotherapy increased activity of VTA GABA neurons in cocaine-experienced rats (Fig 5), we hypothesized that this cell type-specific response would coincide with decreased VTA dopamine cell activity and represent one mechanism by which systemic exendin-4 suppresses cocaine-seeking behavior. We used *in vivo* fiber photometry in transgenic rats to determine how GLP-1R agonist pharmacotherapy alters activity of VTA dopamine neurons during cocaine reinstatement test sessions. To measure changes in intra-cellular calcium levels in midbrain dopamine neurons during cocaine seeking, a Cre-dependent GCaMP8f virus was infused into the VTA of rats expressing Cre-recombinase under the *Th* promoter and a fiberoptic cannula was implanted directly above the infusion site (Fig 6A). A representative image of subregion-specific viral spread is shown in Figure 6B. Fluorescent labelling of *eGfp, Th,* and *Gad1* mRNA transcripts using FISH confirmed selective GCaMP8f expression in *Th*-positive neurons in the VTA (Fig 6B).

**Figure 6.**
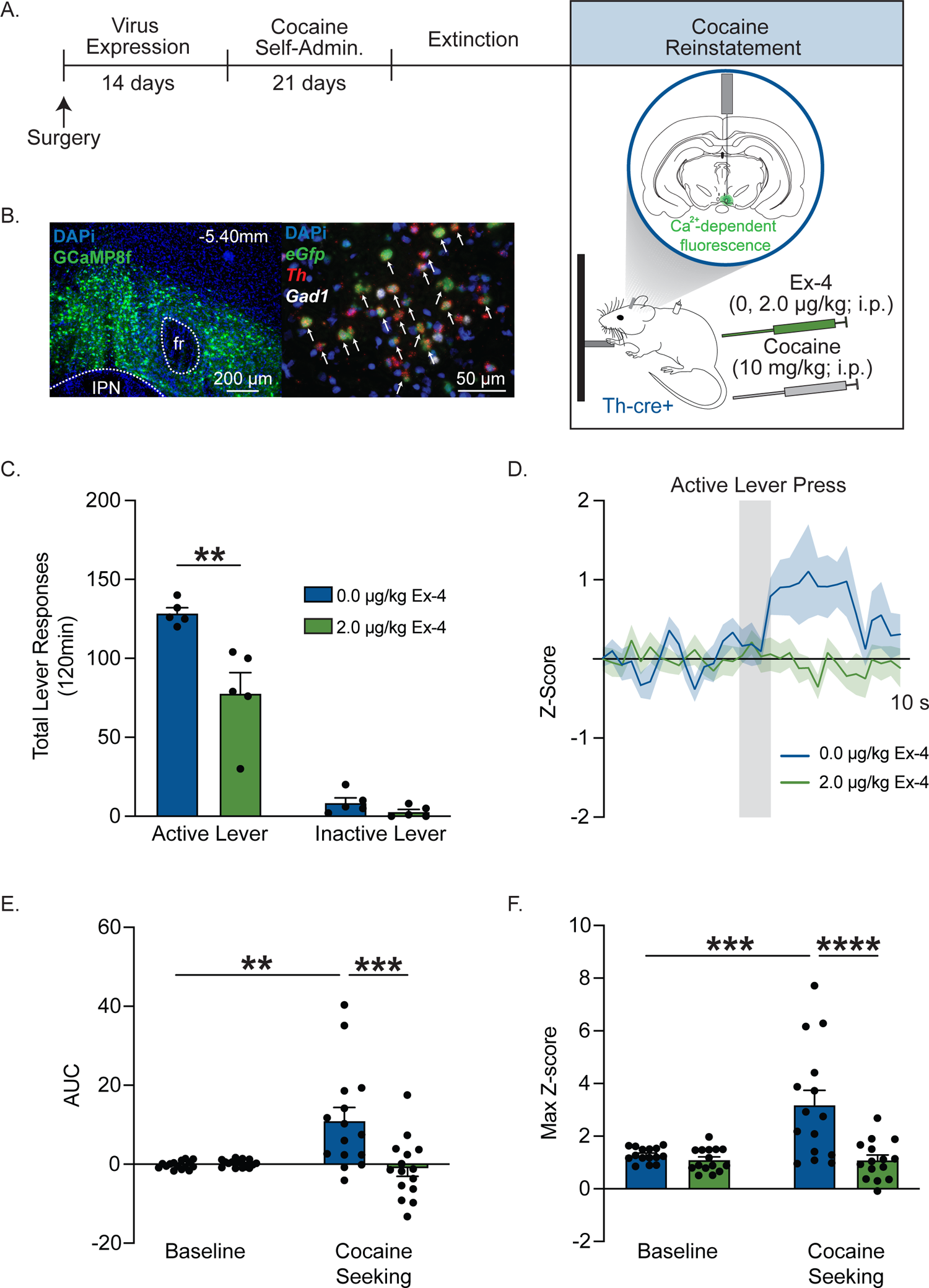
Systemic GLP-1R agonist pharmacotherapy attenuates VTA dopamine neuron activity and decreases cocaine seeking. A. Schematic illustrating viral approach and experiment timeline where a Cre-dependent AAV expressing GCaMP8f virus was infused into the VTA of TH-Cre rats to measure intra-cellular calcium dynamics in VTA dopamine neurons during reinstatement tests sessions. **B** Representative image of GCaMP8f viral expression in the VTA. FISH confirmed selective GCaMP8f expression in *Th*-positive cells [DAPi (blue), *eGfp* (green), *Th* (red), and *Gad1* (white)]. **C** Systemic exendin-4 administration decreased active lever responses during cocaine reinstatement test sessions *(n =* 5; two-way RM ANOVA, treatment x lever: F_1_,_4_ = 39.66, *p* = 0.0032; Bonferroni’s test: vehicle vs exendin-4: \*\**p* = 0.0023). **D** Normalized z-score traces during cocaine seeking in rats pretreated with vehicle or exendin-4 *(n* = 5 rats: 3 active lever presses/rat/treatment). **E** Cocaine seeking significantly increased the AUC of recorded Ca^2+^ signals from VTA dopamine neurons in rats treated with vehicle (two-way RM ANOVA, treatment x time: F_1_,_28_ = 10.71, *p* = 0.0028; Bonferroni’s test: baseline vs cocaine seeking in vehicle-treated controls: ** *p* = 0.0011). Exendin-4 pretreatment significantly attenuated cocaine seeking-evoked increases in VTA dopamine neuron activity (Bonferroni’s test: 0.0 vs 2.0 μg/kg exendin-4 during cocaine seeking: *** *p* = 0.0002). **F** Cocaine seeking significantly increased maximum z-scores in VTA dopamine neurons in rats treated with vehicle (two-way RM ANOVA, treatment x time: F_1_,_28_ = 11.37, *p* = 0.0022; Bonferroni’s test: baseline vs cocaine seeking in vehicle-treated controls: *** *p* = 0.0002). Exendin-4 pretreatment significantly decreased maximum z-scores in VTA dopamine neurons during cocaine seeking compared to vehicle (Bonferroni’s test: cocaine seeking 0.0 μg/kg vs cocaine seeking 2.0 μg/kg, **** *p* < 0.0001). Data are mean ± SEM.

To determine how GLP-1R activation alters calcium dynamics in VTA dopamine neurons during cocaine seeking, rats were pretreated with vehicle or exendin-4 (2.0 µg/kg, i.p.) prior to a cocaine priming-induced reinstatement test session (Fig 6A). Consistent with our prior studies^8,18^ and the reinstatement studies above in GAD-Cre rats (Fig 5C), exendin-4 significantly reduced cocaine seeking in TH-Cre rats (Fig 6C). Cell type-specific normalized z-score traces during cocaine seeking are presented in Figure 6D. AUC and maximum z-scores were significantly increased during cocaine seeking compared to baseline in VTA dopamine neurons of vehicle-treated rats. These findings indicate that cocaine-seeking behavior evokes transient increases in VTA dopamine neuron activity (Fig 6E & 6F). Interestingly, pretreatment with exendin-4 blocked cocaine seeking-evoked increases in AUC and maximum z-scores (Fig 6E & 6F). Normalized z-score traces associated with inactive lever presses during reinstatement tests are presented in Supplementary Figure 5A. No effects of treatment were found on AUC and maximum z-scores from VTA dopamine neurons following inactive lever responses during the reinstatement test sessions (Suppl. Fig 5B & 5C). To confirm that these calcium dynamics were specific to cocaine-seeking behavior, a separate group of TH-Cre rats was allowed to self-administer saline before undergoing extinction and subsequent reinstatement test sessions. There were no effects of vehicle or exendin-4 on AUC or maximum z-scores from VTA dopamine neurons following an active lever press in drug-naïve rats (Suppl. Fig 6A-D). Together, these findings indicate that activity of VTA dopamine neurons is significantly increased during cocaine seeking and that GLP-1R agonist pharmacotherapy prevents/blocks this cellular response and suppresses drug-seeking during withdrawal.

## Discussion

An emerging literature indicates that activating central GLP-1Rs is sufficient to reduce cocaine-seeking behavior^8,14,18^, findings that support translational studies focused on re-purposing GLP-1R agonists to treat CUD^5,22^. However, the neural mechanisms underlying the efficacy of GLP-1R agonists to reduce cocaine seeking and the role of central GLP-1-producing neural circuits in cocaine seeking are unknown. Here, we discovered that cocaine self-administration and acute withdrawal decreased plasma GLP-1 levels and that chemogenetic activation of GLP-1-producing NTS neurons that project to the VTA is sufficient to attenuate cocaine seeking during abstinence. We also showed, for the first time, that GLP-1Rs are expressed primarily on GABA neurons in the VTA and that the suppressive effects of a GLP-1R agonist on cocaine seeking are associated with increased activity of VTA GABA neurons and decreased activity of VTA dopamine neurons. These cell type-specific cellular responses expand our understanding of the neural mechanisms underlying the efficacy of GLP-1R agonists on cocaine seeking. Our findings suggest that targeting endogenous GLP-1-producing neural circuits may reduce cocaine craving-induced relapse and further support the potential for GLP-1R-based pharmacotherapeutic approaches to treat CUD.

### Endogenous GLP-1 signaling is dynamically altered by cocaine taking and abstinence

A pilot study of human cocaine users showed that intravenous cocaine self-administration decreased serum GLP-1 levels, effects associated with cocaine-related cardiorespiratory and subjective responses^35^. While it is not clear how decreased serum GLP-1 may promote the subjective experiences of cocaine in humans, these results suggest that reduced GLP-1 signaling may drive on-going cocaine taking. We extended these findings to a preclinical model of CUD and found that plasma GLP-1 concentrations were significantly decreased following intravenous cocaine self-administration and acute abstinence in rats. Given that systemic administration of GLP-1R agonists reduce cocaine taking^36^ and seeking^8,18^ in rodents, these results suggest that decreased endogenous GLP-1 signaling may facilitate cocaine-seeking behavior. Consistent with these results, expression of PPG mRNA, a necessary precursor of GLP-1 production, is decreased in the NTS following seven days of cocaine abstinence, a time point associated with robust cocaine-seeking behavior in rats^8^. Together, these findings support the hypothesis that decreased endogenous GLP-1 signaling promotes/facilitates drug seeking and that activating central GLP-1-producing neural circuits may attenuate cocaine-seeking behavior during abstinence.

### Targeted activation of endogenous GLP-1-producing ‘anti-craving’ circuits attenuates cocaine seeking

The VTA receives monosynaptic projections from GLP-1-producing neurons in the NTS^37^ and plays a critical role in the reinstatement of cocaine-seeking behavior^4^. While cocaine taking does not alter GLP-1R expression in the VTA, abstinence following cocaine self-administration decreased PPG mRNA expression in the NTS^8^. These results suggest that decreased PPG expression in the NTS may facilitate/promote cocaine seeking via reduced GLP-1 tone in target nuclei, including the VTA^5^. Here, we showed that chemogenetic activation of GLP-1-producing NTS neurons that project to the VTA is sufficient to suppress drug seeking in cocaine-experienced rats. These findings are the first to establish a functional role for GLP-1-producing NTS→VTA projections in cocaine seeking and support the hypothesis that endogenous GLP-1R-expressing circuits are important ‘anti-craving’ pathways that when activated reduce drug seeking during abstinence. These results are consistent with a previous study in which we reported that chemogenetic activation of GLP-1-producing NTS neurons that project to the laterodorsal tegmental nucleus (LDTg) reduced cocaine seeking^14^. These effects were mediated, in part, by activation of GLP-1Rs on GABAergic LDTg neurons that project to the VTA^14^. Collectively, these findings indicate that activation of GLP-1-producing neurons in the NTS attenuates cocaine seeking and regulates the mesolimbic dopamine system via direct projections to the VTA as well as polysynaptic connections to the VTA via GLP-1R-expressing hubs such as the LDTg.

Previous studies in drug-naïve rodents showed that activation of GLP-1-producing NTS neurons that project to the VTA transiently reduced food intake^28^. Consistent with these effects, systemic infusions of GLP-1R agonists attenuate food intake in drug-naïve rodents and humans^20,30,38^. Therefore, we screened for these potential adverse effects following NTS→VTA circuit activation in our cocaine-experienced animals. We found that activation of NTS→VTA circuits suppressed cocaine seeking in rats without affecting *ad libitum* food and water intake. Thus, targeted activation of endogenous GLP-1-producing neural circuits selectively reduces drug seeking during abstinence in cocaine-experienced rats. Although it is not clear how these findings will translate to humans, these preclinical studies indicate that activation of NTS→VTA GLP-1-producing projections selectively reduce cocaine seeking. In addition to monitoring weight loss and nutritional statues, future clinical trials should measure nausea and emesis following activation of central GLP-1-producing circuits as these adverse effects are the main reason cited for discontinuation of GLP-1R agonist pharmacotherapy^29,39^.

To date, only one other study has investigated the functional role of NTS→VTA circuits in motivated behaviors. This study showed that chemogenetic activation of NTS→VTA projections decreased consumption of a high fat diet (HFD) in drug-naïve mice^28^. In contrast, our findings clearly show that chemogenetic activation of NTS→VTA projections during abstinence did not alter *ad libitum* food intake or body weight in cocaine-experienced rats. These findings are consistent with our previous behavioral pharmacology studies in which we identified behaviorally selective doses of systemic and intra-VTA exendin-4 that reduced drug seeking in cocaine-experienced rats and did not alter food intake or body weight^8^. Although not clear, these discrepant findings could be due to species differences, cocaine exposure, and/or selective effects of NTS→VTA circuit activation on macronutrient preference (HFD vs. normal chow). In support of the latter, chemogenetic activation of the NTS suppressed consumption of a HFD but did not alter intake of normal chow in drug-naïve mice^28^. A limitation of this study was the activation of heterogenous NTS neuron populations (i.e., Neither GLP-1-producing cells or projection-specific neurons were targeted). It is possible that additional satiety signals, such as glutamate^40^, GABA^41^, cholecystokinin (CCK)^42^, pro-opiomelanocortin (POMC)^43^, neuropeptide Y^44^, and/or norepinephrine^44^, are mediating the macronutrient-dependent feeding responses associated with NTS neuron activation. However, given that GLP-1R agonism in the VTA more robustly suppresses the intake of highly palatable foods compared to normal chow in drug-naïve rodents^37,45,46^, it is likely that endogenous GLP-1 signaling in the VTA more selectively regulates hedonic rewards versus homeostatic signals^4^. Our preclinical evidence to date indicates that cocaine-mediated behaviors are more sensitive to modulation by GLP-1R agonists and increased central GLP-1 signaling than non-drug motivated behaviors in cocaine-experienced rats.

### The efficacy of systemic GLP-1R agonists to attenuate cocaine seeking is associated with increased activity of GABA neurons in the VTA

While an emerging literature indicates that activation of VTA GLP-1Rs is sufficient to reduce drug-seeking behavior^8,14^, the cell type-specific neural mechanisms underlying the efficacy of GLP-1R agonists and increased GLP-1 signaling in the VTA on cocaine seeking remain largely unknown. Using FISH and snRNA-seq, we discovered that GLP-1Rs are expressed primarily on GABA neurons in the rat VTA. No *Glp1r* transcripts were identified in VTA dopamine neurons. This anatomy suggests that enhanced GLP-1 signaling in the VTA regulates cocaine seeking, in part, through activation of GLP-1Rs expressed on inhibitory midbrain circuits. These data are consistent with previous studies of transgenic mice modified to express fluorescent proteins under the *Glp1r* promoter^47,48^. These studies showed little to no co-expression GLP-1Rs and dopamine cell markers in the VTA^47,48^ and high co-expression of GLP-1Rs and GABA cell markers throughout the mesolimbic reward system^47^. We also identified populations of *Glp1r*-expressing cells in the VTA that did not express *Gad1 or Th*. Future studies are needed to characterize these cell populations and determine their functional relevance to cocaine-mediated behaviors.

GLP-1Rs are coupled predominantly to G_s_-proteins and increase cAMP signaling when activated^49,50^. Electrophysiology studies in drug-naïve rodents revealed that GLP-1R agonism transiently increased the frequency and amplitude of spontaneous and miniature inhibitory post-synaptic currents^51,52,53^. Together with our present findings that GLP-1Rs are expressed primarily on VTA GABA neurons and that GLP-1R agonism increases c-Fos expression in VTA neurons, we hypothesized that systemic administration of a GLP-1R agonist would increase the activity of VTA GABA neurons and decrease cocaine seeking. Here, we showed that exendin-4 increased VTA GABA neuron activity during cocaine seeking at a dose that significantly attenuated cocaine reinstatement. These findings highlight a novel mechanism wherein GLP-1R agonist pharmacotherapy activates inhibitory midbrain neurons to suppress cocaine-seeking behavior. These data are also consistent with prior studies in which stimulation of GABA neurons in the VTA with GABA receptor agonists attenuated the ability of cocaine^54^, cocaine-paired cues^55^, and stress^56^ to reinstate cocaine-seeking behavior. Importantly, the current study is limited to the effects of GLP-1R activation on VTA GABA neuron activity during cocaine-primed reinstatement. Future studies are needed to determine if GLP-1R pharmacotherapy has similar effects on VTA GABA neurons during cue-induced and/or stress-induced reinstatement. It is possible that the GLP-1R-expressing circuits that mediate the ability of these distinct stimuli to reinstate cocaine-seeking behavior are different. Given the high co-expression of GLP-1Rs and GABA cell markers throughout the mesolimbic system^14,47^, GABA neurons in other nuclei implicated in cocaine seeking are anatomically and functionally positioned to contribute to the efficacy of GLP-1R pharmacotherapy. Therefore, future studies should also explore how GLP-1R pharmacotherapy influences the activity of GABA neurons in other relevant nuclei, such as the LDTg, the central nucleus of the amygdala (CeA), and the nucleus accumbens (NAc), as well as how these GLP-1R-expressing GABA circuits regulate cocaine-mediated behaviors.

Previously, we showed that systemic exendin-4 penetrates the brain and distributes to the VTA where it is localized in proximity to *Th+* neurons^8^. Anatomical studies, albeit in mice, identified dense GLP-1R-expressing fibers surrounding VTA dopamine neurons^47^. These data suggest a presynaptic mechanism by which GLP-1R-expressing terminals are modulating phasic dopamine cell firing in the VTA. Indeed, activation of GLP-1Rs expressed on presynaptic terminals in the hippocampus has been shown to enhance depolarization-evoked GABA release^57^. It is possible that GLP-1R agonists reduce cocaine seeking, in part, by increasing GABA release from presynaptic terminals in the VTA, effects consistent with the post-synaptic mechanism revealed in the current study. In addition, one study suggested that activating presynaptic GLP-1Rs may alter glutamate signaling in the VTA and, in turn, modulate midbrain dopamine cell activity^45^. Thus, the neurochemical mechanisms in the midbrain underlying the efficacy of GLP-1R agonists on cocaine seeking are complex and likely involve both presynaptic and postsynaptic GLP-1Rs. Future studies are needed to fully characterize the role of presynaptic VTA GLP-1Rs in modulating phasic dopamine cell firing and cocaine-seeking behavior.

### The efficacy of systemic GLP-1R agonists to attenuate cocaine seeking is associated with decreased activity of dopamine cells in the VTA

VTA dopamine neurons mediate the rewarding effects of cocaine and increased dopamine cell firing in the VTA is necessary for cocaine seeking^58,59^. For example, chemogenetic inhibition of VTA dopamine neurons prevented the ability of a priming injection of cocaine and stress to reinstate drug seeking in cocaine-experienced rats^59^. Additionally, *in vivo* electrophysiology studies in awake, freely behaving rats have shown that cocaine increases VTA dopamine cell-firing^60,61^ and cocaine self-administration produces long-term changes in VTA dopamine neuron excitability to drive cocaine seeking^62^. In the current study, we showed that exendin-4 attenuates cocaine seeking and blocks cocaine seeking-evoked increases in VTA dopamine activity. These findings are consistent with microdialysis studies showing that systemic doses of exendin-4 that attenuated cocaine-induced conditioned place preference and cocaine self-administration in mice also decreased cocaine-evoked dopamine release in the NAc^32^ and dorsal striatum^36^. Similarly, a study done in head-fixed rats using fast-scan cyclic voltammetry showed that exendin-4 pretreatment suppressed the induction of phasic dopamine release events in the NAc by intravenous cocaine^63^. Collectively, these findings suggest that increased GLP-1R activation in the VTA attenuates cocaine seeking, in part, by decreasing

phasic dopamine cell firing. Future studies are needed to explore how GLP-1R agonism influences dopamine release in the striatum, as well as the activity of D1 dopamine receptor- and D2 dopamine receptor-expressing medium spiny neurons in the NAc during cocaine seeking. These studies will provide important insights into the how GLP-1R activation in the VTA alters dopamine signaling to influence activity in the two main output pathways of the striatal complex and ultimately decrease cocaine seeking.

Comprehensively, our results indicate that GLP-1R agonist pharmacotherapy produces distinct cellular responses in VTA GABA and dopamine neurons of cocaine-experienced rats. Specifically, we found GLP-1R agonist pharmacotherapy increased VTA GABA neuron activity associated with cocaine seeking and suppressed cocaine-seeking evoked increases in VTA dopamine neuron activity. Importantly, there are heterogenous populations of GABA neurons in the VTA, including GABA interneurons. VTA GABA interneurons provide a local source of inhibitory control^64^ and disrupt reward-mediated behaviors^65,66,67^. Thus, our data suggest a novel neural mechanism by which GLP-1R agonists attenuate cocaine seeking through stimulation of local GABA neurons that suppress VTA dopamine cell firing (Fig 7). In addition to GABA interneurons, we hypothesize that activation of presynaptic GLP-1Rs promotes GABA release onto VTA dopamine neurons to suppress their activity during cocaine seeking. Since VTA GABA neurons in the VTA have also been shown to be outwardly projecting, it is also possible that GLP-1Rs are expressed on GABA neurons in the VTA that project to downstream nuclei including the NAc, CeA, prefrontal cortex, lateral hypothalamus, and the lateral habenula^68,69,70^ . Future studies are needed to comprehensively characterize the anatomy of GLP-1R-expressing inhibitory midbrain circuits and their unique functional roles in cocaine-seeking behavior.

**Figure 7.**
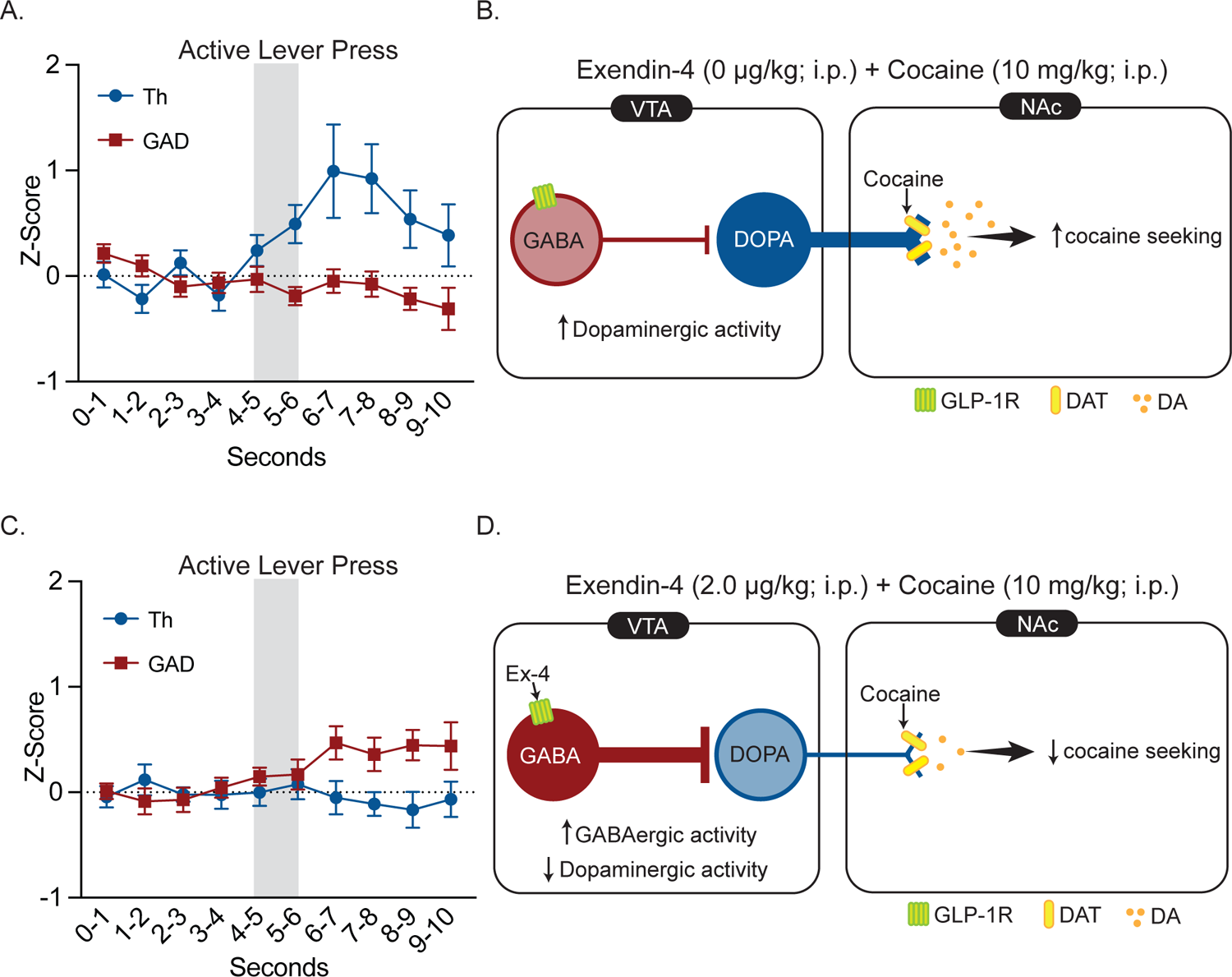
GLP-1R agonist pharmacotherapy attenuates cocaine seeking by activating inhibitory VTA GABA neurons and decreasing activity of VTA dopamine neurons. A Normalized and down-sampled dopamine and GABA z-score traces during cocaine seeking from rats pretreated with vehicle prior to cocaine priming-induced reinstatement test sessions. **B** Schematic illustrating cocaine seeking-evoked increases in dopamine cell firing in the VTA. **C** Normalized and down-sampled dopamine and GABA z-score traces during cocaine seeking from rats pretreated with exendin-4 prior to cocaine priming-induced reinstatement test sessions. **D** Schematic illustrating one proposed mechanism by which exendin-4 increases VTA GABA cell firing which, in turn, inhibits VTA dopamine cell firing and suppresses cocaine-seeking behavior. Additional mechanisms likely include regulation of presynaptic GABA and/or glutamate release (see main text for more details).

## Conclusion

Here, we identified a novel endogenous GLP-1-producing neural circuit that regulates cocaine-seeking behavior. We also discovered a novel cell type-specific mechanism in the VTA underlying the efficacy of a GLP-1R agonist on cocaine seeking. These data highlight a potential therapeutic approach to targeting endogenous GLP-1-producing ‘anti-craving’ circuits to reduce cocaine relapse and further support of the translational potential of GLP-1R agonists as pharmacotherapies for CUD. Our findings also inform innovative strategies in the development of next-generation GLP-1R-based therapies to treat CUD. Studies done in drug-naïve animals revealed that activation GLP-1-producing neurons in the NTS are not necessary for the anorexic effects of systemic GLP-1R agonists and that concurrent NTS GLP-1-producing circuit activation suppresses feeding more potently than GLP-1R agonism alone^71^. Therefore, simultaneous activation of endogenous GLP-1-producing ‘anti-craving’ circuits and systemic GLP-1R agonist administration is a provocative therapeutic strategy that may suppress drug seeking to a greater degree than either intervention alone in cocaine-experienced rats.

## Acknowledgements

The authors would like to thank Jennifer Ben Nathan, Suraj Neelamagam, and Antonia Caffrey for their technical contributions to this project.

## Author contributions

RM, NH, and VW contributed to the acquisition and analyses of the data as well as drafted the manuscript. VW, YZ, MTR, RCC, and BCR, contributed to data collection and analyses. HDS was responsible for the study concept and design, supervised the acquisition of the data, and helped draft the manuscript. All authors participated in reviewing and editing the manuscript for publication.

