## Supplementary Materials for "An endogenous GLP-1 circuit engages VTA GABA neurons to regulate mesolimbic dopamine neurons and attenuate cocaine seeking"

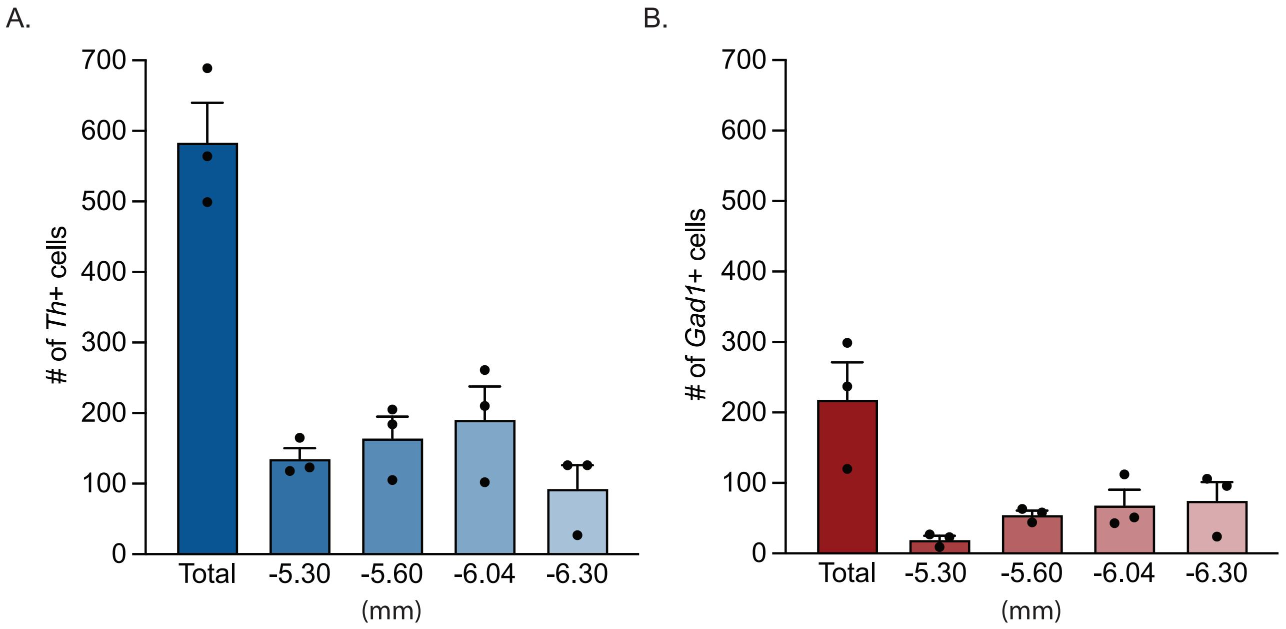


**Supplementary Figure 1. Distribution of *Th*+ and *Gad1*+ cells in the VTA.** The number of *Th*+ cells (**A**) and *Gad1*+ cells (**B**) in the VTA at different anterior/posterior positions relative to bregma ($n$ = 3 rats; 4 slices per rat). Data are mean ± SEM.

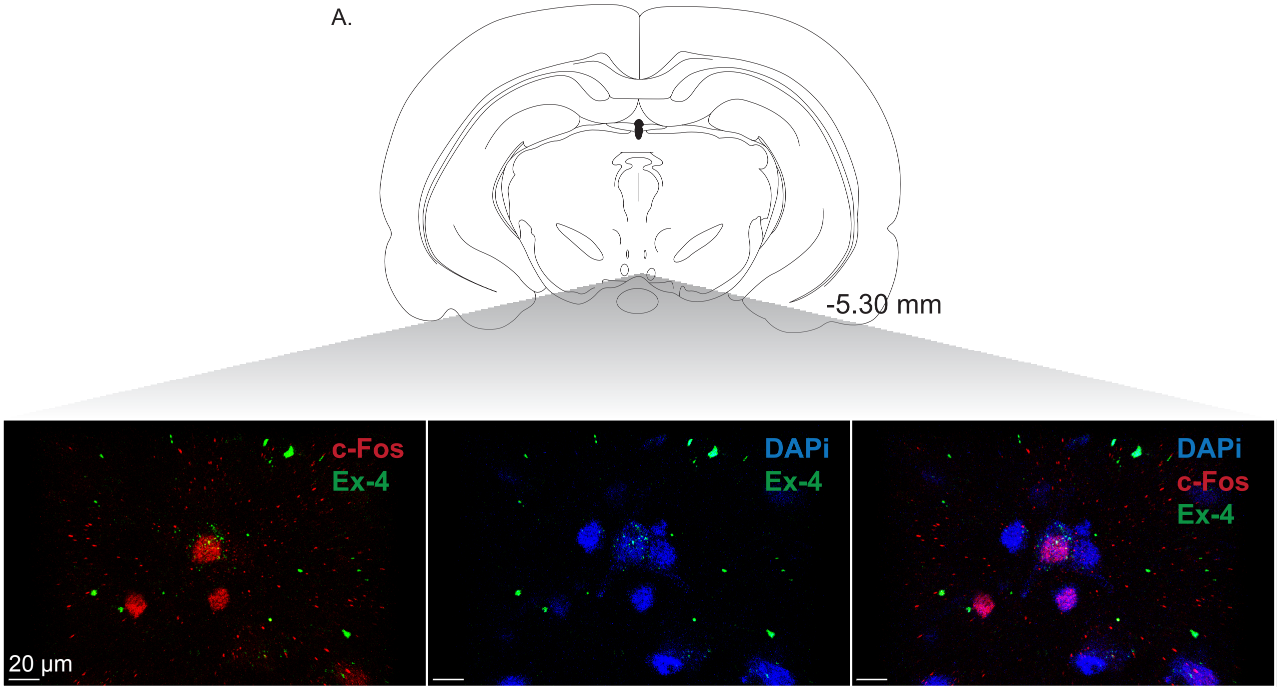


**Supplementary Figure 2. Systemically administered fluoro-exendin-4 binds to putative GLP-1Rs on neurons in the VTA and induces c-Fos expression.** A drug-naïve rat was pretreated with 0.2 µg/kg fluoro-exendin-4 (i.p.) and then sacrificed 90 minutes later. Immunohistochemistry identified fluoro-exendin-4 bound to VTA neurons expressing c-Fos [DAPi (blue), exendin-4 (green), and c-Fos (red)] (*n* = 1). Data are mean ± SEM.

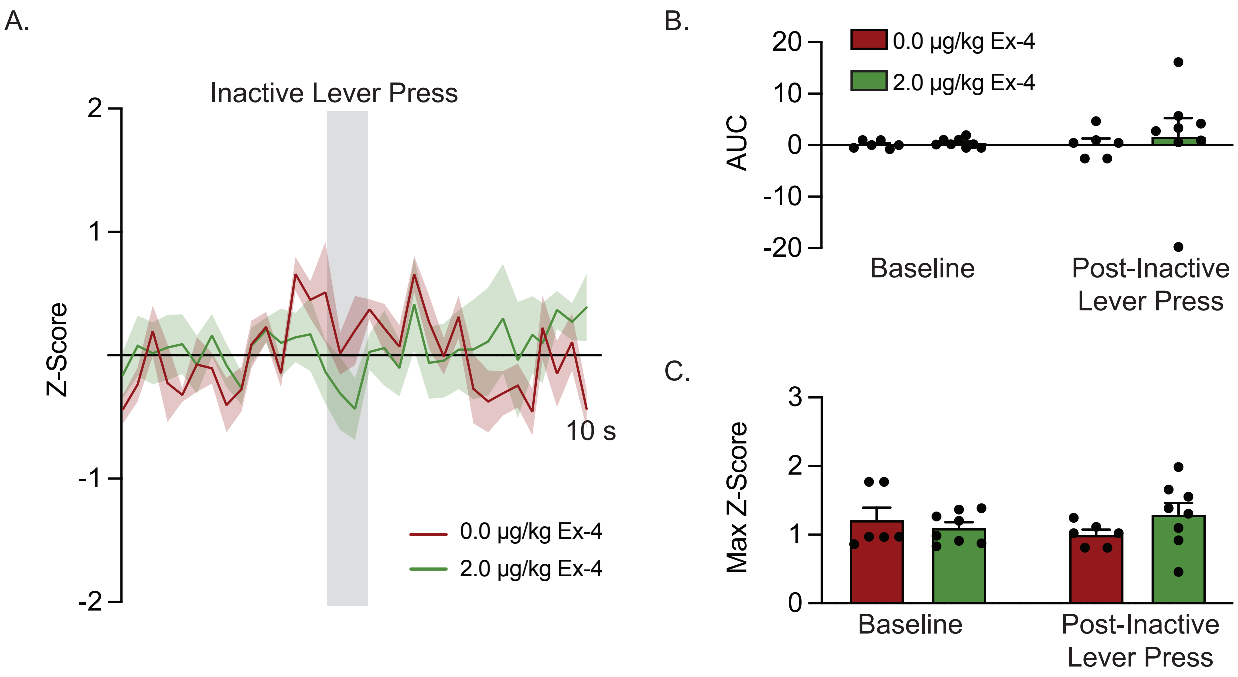


**Supplementary Figure 3. Systemic GLP-1R agonist pharmacotherapy does not alter VTA GABA neuron activity associated with inactive lever presses during cocaine reinstatement test sessions. A** Normalized z-score traces from inactive lever presses during reinstatement test sessions in cocaine-experienced rats pretreated with vehicle or exendin-4 ($n$ = 5 rats; 2 inactive lever presses/rat/treatment). Some rats did not press the inactive lever during reinstatement tests (0.0 µg/kg exendin-4: $n$ = 6 presses from 3 rats; 2.0 µg/kg exendin-4: $n$ = 8 presses from 4 rats). **B** Responding on the inactive lever during reinstatement tests had no effect on the AUC of recorded $Ca^{2+}$ signals from VTA GABA neurons in cocaine-experienced rats treated with vehicle or exendin-4 (two-way ANOVA, treatment: $F_{1,24}$ = 0.1768, $p$ = 0.6778, time: $F_{1,24}$ = 0.1082, $p$ = 0.7450, treatment x time: $F_{1,24}$ = 0.07757, $p$ = 0.7830). **C** Responding on the inactive lever during reinstatement tests had no effect on maximum z-scores from VTA GABA neurons in cocaine-experienced rats treated with vehicle or exendin-4 (two-way ANOVA, treatment: $F_{1,24}$ = 0.4365, $p$ = 0.5151, time: $F_{1,24}$ = 0.005857, $p$ = 0.9396, treatment x time: $F_{1,24}$ =0.1452, $p$ = 0.1452). Data are mean ± SEM.

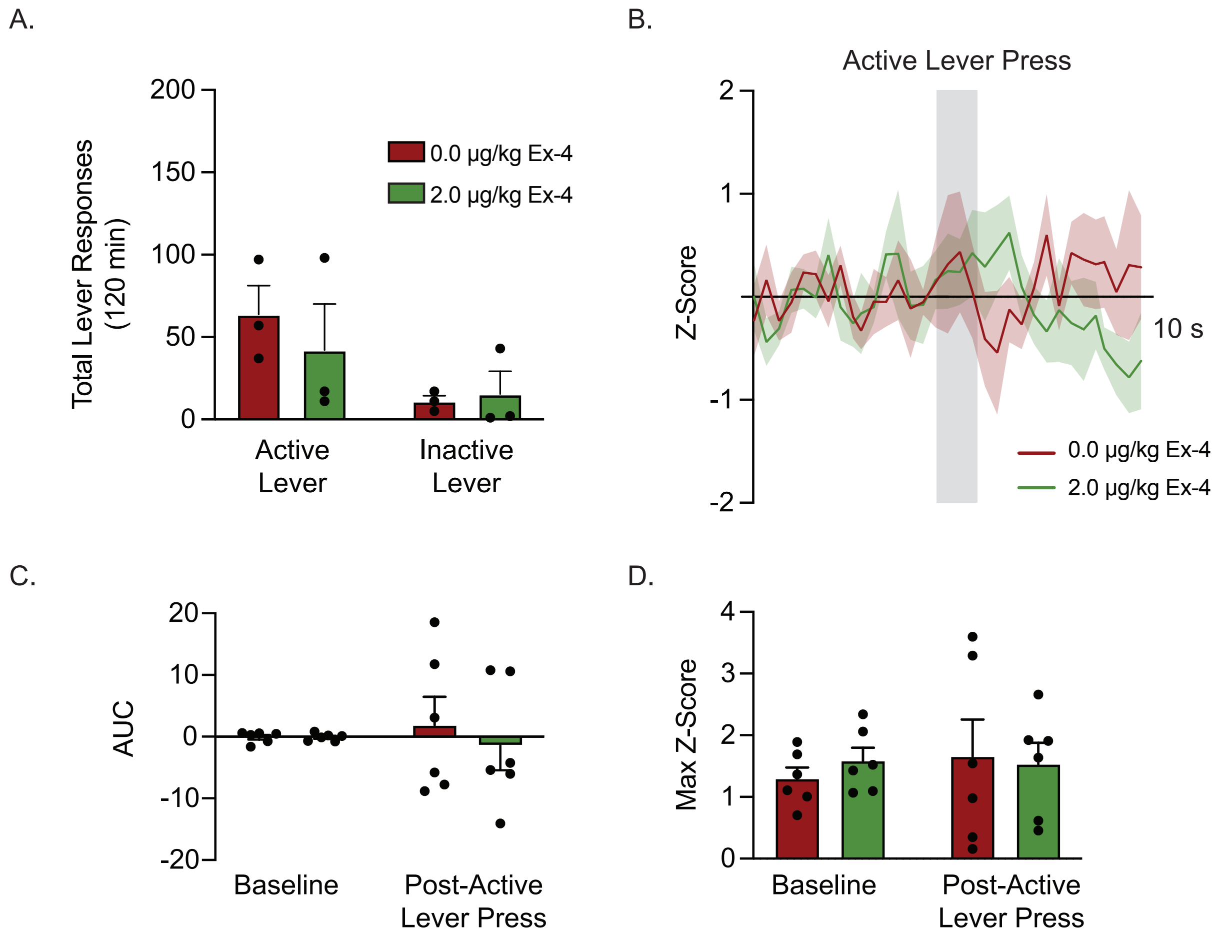


**Supplementary Figure 4. Systemic GLP-1R agonist administration does not alter VTA GABA neuron activity associated with active lever presses in drug-naïve rats. A** Systemic exendin-4 administration had no effect on total lever responses during reinstatement test sessions in rats that previously self-administered saline (*n* = 3; two-way RM ANOVA, treatment: treatment: $F_{1,2}$ = 6.969, *p* = 0.1185, lever: $F_{1,2}$ = 2.648, *p* = 0.2452, treatment x time: $F_{1,2}$ = 0.7533, *p* = 0.4769). **B** Normalized z-score traces during active lever presses in control rats pretreated with vehicle or exendin-4 *(n* = 3 rats; 2 active lever presses/rat/treatment). **C** Responding on the active lever had no effect on the AUC of recorded $Ca^{2+}$ signals from VTA GABA neurons in drug-naïve rats treated with vehicle or exendin-4 (*n* = 6; two-way RM ANOVA, treatment: $F_{1,10}$ = 0.2900, *p* = 0.6020, time: $F_{1,10}$ = 0.008872, *p* = 0.9268, treatment x time: $F_{1,10}$ = 0.2568, *p* = 0.6233). **D** Responding on the active lever had no effect on maximum z-scores from VTA GABA neurons in drug-naïve rats treated with vehicle or exendin-4 (two-way RM ANOVA, treatment: $F_{1,10}$ = 0.06496, *p* = 0.8040, time: $F_{1,10}$ = 0.1378, *p* = 0.7182, treatment x time: $F_{1,10}$ = 0.2498, *p* = 0.6281). Data are mean ± SEM.

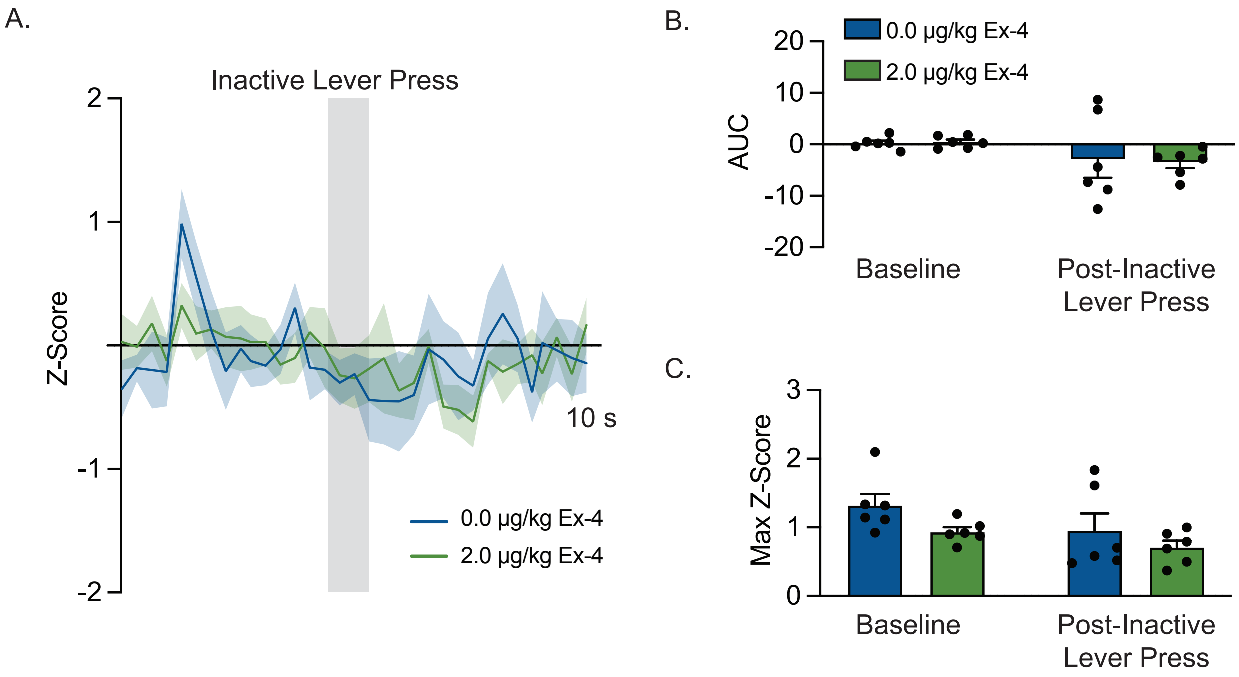


**Supplementary Figure 5. Systemic GLP-1R agonist pharmacotherapy does not alter VTA dopamine neuron activity associated with inactive lever presses during cocaine reinstatement test sessions. A** Normalized z-score traces from inactive lever presses during reinstatement test sessions in cocaine-experienced rats pretreated with vehicle or exendin-4 (*n* = 5 rats; 2 inactive lever presses/rat/treatment). Some rats did not press the inactive lever during recording (0.0 µg/kg exendin-4: *n* = 6 presses from 3 rats; 2.0 µg/kg exendin-4: *n* = 6 presses from 3 rats). **B** Responding on the inactive lever during reinstatement tests had no effect on the AUC of recorded $Ca^{2+}$ signals from VTA dopamine neurons in cocaine-experienced rats treated with vehicle or exendin-4 (two-way ANOVA, treatment: $F_{1,20}$ = 0.1161, *p* = 0.9153, time: $F_{1,20}$ = 3.635, *p* = 0.0710, treatment x time: $F_{1,20}$ = 0.04728, *p* = 0.8301). **C** Responding on the inactive lever during reinstatement tests had no effect on maximum z-scores from VTA GABA neurons in cocaine-experienced rats treated with vehicle or exendin-4 (two-way ANOVA, treatment: $F_{1,20}$ = 3.891, *p* = 0.0625, time: $F_{1,20}$ = 3.413, *p* = 0.0795, treatment x time: $F_{1,20}$ = 0.1941, *p* = 0.6642). Data are mean ± SEM.

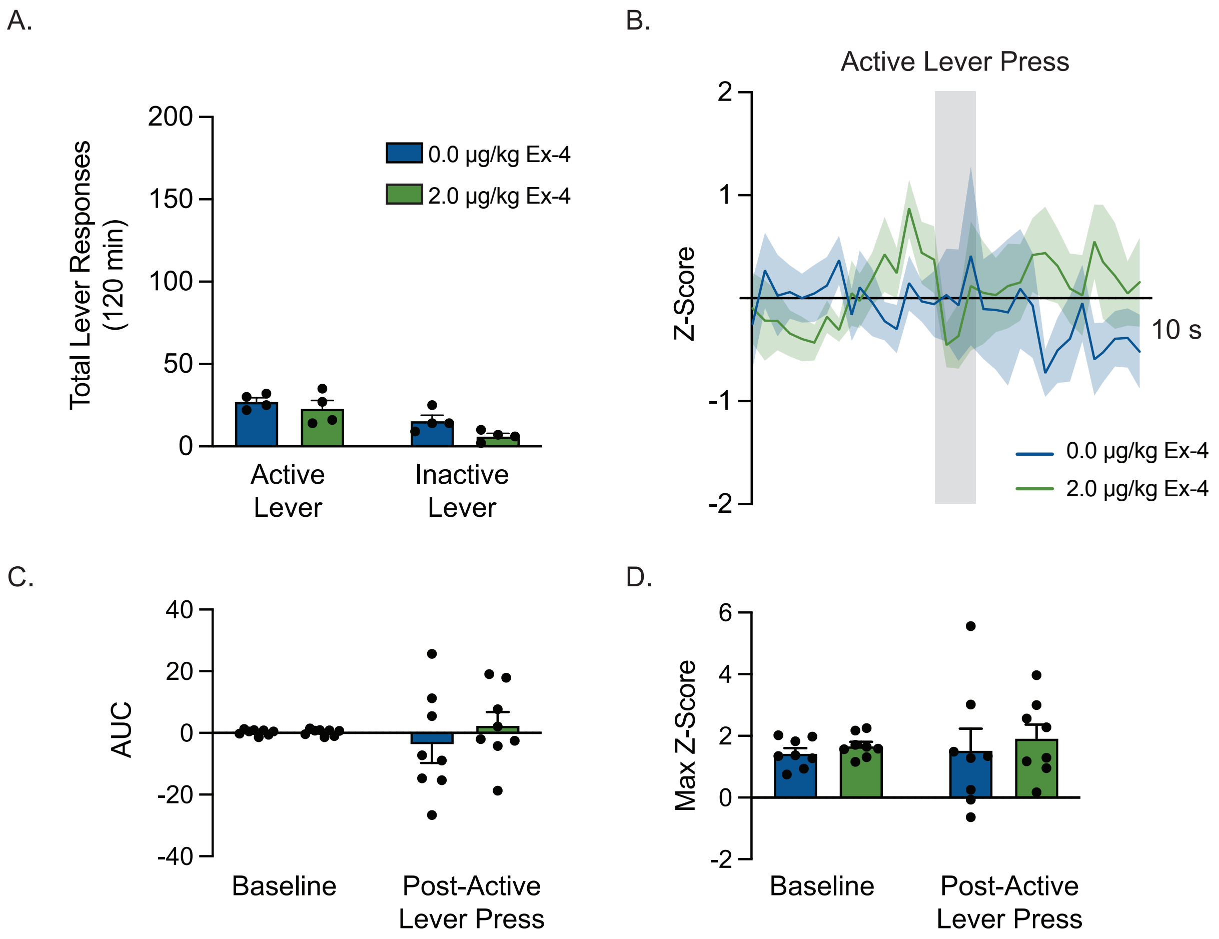


**Supplementary Figure 6. Systemic GLP-1R agonist administration does not alter VTA dopamine neuron activity associated with active lever presses in drug-naïve rats. A** Systemic exendin-4 administration had no effect on total lever responses during reinstatement test sessions in rats that previously self-administered saline (*n* = 4; two-way RM ANOVA, treatment: treatment: $F_{1,3}$ = 5.622, *p* = 0.0984, lever: $F_{1,3}$ = 9.976, *p* = 0.0509, treatment x time: $F_{1,2}$ = 0.8876, *p* = 0.4156). **B** Normalized z-score traces during active lever presses in control rats pretreated with vehicle or exendin-4 *(n* = 4 rats; 2 active lever presses/rat/treatment). **C** Responding on the active lever had no effect on the AUC of recorded $Ca^{2+}$ signals from VTA dopamine neurons in drug-naïve rats treated with vehicle or exendin-4 (*n* = 8; two-way RM ANOVA, treatment: $F_{1,14}$ = 0.7561, *p* = 0.3992, time: $F_{1,14}$ = 0.05920, *p* = 0.8113, treatment x time: $F_{1,14}$ = 0.6591, *p* = 0.4305). **D** Responding on the active lever had no effect on maximum z-scores from VTA dopamine neurons in drug-naïve rats treated with vehicle or exendin-4 (two-way RM ANOVA, treatment: $F_{1,14}$ = 0.1463, *p* = 0.7078, time: $F_{1,14}$ = 0.6045, *p* = 0.4498, treatment x time: $F_{1,14}$ = 0.03093, *p* = 0.8629). Data are mean ± SEM.
